# Frequent loss-of-heterozygosity in CRISPR-Cas9-edited early human embryos

**DOI:** 10.1101/2020.06.05.135913

**Authors:** Gregorio Alanis-Lobato, Jasmin Zohren, Afshan McCarthy, Norah M.E. Fogarty, Nada Kubikova, Emily Hardman, Maria Greco, Dagan Wells, James M.A. Turner, Kathy K. Niakan

**Author notes:** To whom correspondence should be addressed: Kathy K. Niakan, Human Embryo and Stem Cell Laboratory, The Francis Crick Institute, 1 Midland Road London NW1 1AT UK.

## Abstract

CRISPR-Cas9 genome editing is a promising technique for clinical applications, such as the correction of disease-associated alleles in somatic cells. The use of this approach has also been discussed in the context of heritable editing of the human germline. However, studies assessing gene correction in early human embryos report low efficiency of mutation repair, high rates of mosaicism and the possibility of unintended editing outcomes that may have pathologic consequences. We developed computational pipelines to assess single-cell genomics and transcriptomics datasets from OCT4 (*POU5F1*) CRISPR-Cas9-targeted and control human preimplantation embryos. This allowed us to evaluate on-target mutations that would be missed by more conventional genotyping techniques. We observed loss-of-heterozygosity in edited cells that spanned regions beyond the *POU5F1* on-target locus, as well as segmental loss and gain of chromosome 6, on which the *POU5F1* gene is located. Unintended genome editing outcomes were present in approximately 16% of the human embryo cells analysed and spanned 4 to 20kb. Our observations are consistent with recent findings indicating complexity at on-target sites following CRISPR-Cas9 genome editing. Our work underscores the importance of further basic research to assess the safety of genome editing techniques in human embryos, which will inform debates about the potential clinical use of this technology.

## Introduction

Clustered regularly interspaced short palindromic repeat (CRISPR)-CRISPR associated 9 (Cas9) genome editing is not only an indispensable molecular biology technique (1) but also has enormous therapeutic potential as a tool to correct disease-causing mutations (2). Genome editing of human embryos or germ cells to produce heritable changes has the potential to reduce the burden of genetic disease and its use in this context is currently a topic of international discussions centred around ethics, safety and efficiency (3, 4).

Several groups have conducted studies to assess the feasibility of gene correction in early human embryos (5–7) and they all encountered low efficiency of gene repair and high levels of mosaicism (i.e. embryos with corrected as well as mutant uncorrected blastomeres or blastomeres with unintended insertion/deletion mutations), which are unacceptable outcomes for clinical applications. In 2017, Ma *et al.* set out to correct a 4bp pathogenic heterozygous deletion in the *MYBPC3* gene using the CRISPR-Cas9 system (8). The experimental strategy involved co-injection of Cas9 protein, a single guide RNA (sgRNA) that specifically targeted the *MYBPC3* mutation and a repair template into either fertilised eggs (zygotes) or oocytes, coincident with intracytoplasmic sperm injection. Analysis of the resulting embryos revealed a higher than expected incidence, with respect to controls, of samples where only wild-type copies of the gene were detectable (8). Intriguingly, the excess of apparently uniformly homozygous wild-type embryos in both cases was not associated with use of the provided repair template for gene correction. Instead, the authors suggest that in edited embryos the wild-type maternal allele served as a template for the high-fidelity homology directed repair (HDR) pathway to repair the double-strand lesion caused by the Cas9 protein in the paternal allele (8).

Ma and colleagues’ interpretation of gene editing by inter-homologue homologous recombination (IH-HR) in the early human embryo has been met with scepticism because alternative explanations can account for the observed results (9–11). One of these is that the CRISPR-Cas9 system can induce large deletions and complex genomic rearrangements with pathogenic potential at the on-target site (9, 10, 12–14). These events can be overlooked because genotyping of the targeted genomic locus often involves the amplification of a small PCR fragment centred around the on-target cut-site. CRISPR-Cas9-induced deletions larger than these fragments in either direction would eliminate one or both PCR primer annealing sites. This in turn can lead to amplification of only one allele, giving the false impression that targeting was unsuccessful or that there is a single homozygous event at the on-target site (9, 10, 15). Loss-of-heterozygosity (LOH) can also be the result of more complex genomic rearrangements like inversions, large insertions, translocations, chromosome loss and even IH-HR with crossover, whereby a large piece of one parental allele is integrated by the other parental chromosome at the on-target cut-site (15).

The reported frequencies of unintended CRISPR-Cas9 on-target damage are not negligible. Adikusama *et al.* targeted six genes in a total of 127 early mouse embryos and detected large deletions (between 100bp and 2.3kb) in 45% of their samples using long-range PCR (10). Of note, large deletions were generally more prevalent when they targeted intronic regions (>70%) than when they targeted exons (20%). Consistent with this, Kosicki and colleagues observed large deletions (up to 6kb) and other complex genomic lesions at frequencies of 5-20% of their clones after targeting the *PigA* and *Cd9* loci in two mouse embryonic stem cell (mESC) lines and primary mouse cells from the bone marrow, as well as the *PIGA* gene in immortalised human female retinal pigment epithelial cells (12). Moreover, Owens *et al.* used CRISPR-Cas9 with two sgRNAs to delete 100-150bp in the *Runx1* locus of mESCs and found that 23% of their clones had large deletions (up to 2kb) that escaped genotyping by short-range PCR (giving the impression that they were homozygous wild-type clones), with these complex on-target events becoming evident using long-range PCR (14). Similar damage and frequencies were also observed with the Cas9^D10A^ nickase (14). More dramatic events were identified by Cullot *et al.*, who CRISPR-targeted the *UROS* locus in HEK293T and K562 cells for HDR correction with a repair template (13). Their experiments suggest that CRISPR-Cas9 can induce mega-base scale chromosomal truncations (~10% increase compared to controls). However, these cells have abnormal karyotypes and are p53 deficient, which may impact on their DNA damage repair machinery. In fact, they did not see the same effect in human foreskin fibroblasts but knocking-out of *TP53* in these primary cells increased the large deletion events by 10-fold (13). More recently, Przewrocka and colleagues observed a 6% incidence of chromosome arm truncations when targeting *ZNF516* in p53-competent HCT116 cancer cell lines with CRISPR-Cas9, suggesting that TP53 expression alone may not predict predisposition of cells to large on-target mutations (16).

Our laboratory used CRISPR-Cas9 genome editing to investigate the function of the pluripotency factor OCT4 (encoded by the *POU5F1* gene on the p-arm of chromosome 6) during human preimplantation development (17). We generated a number of single-cell amplified genomic DNA (gDNA) samples for genotyping and confirmed on-target genome editing in all microinjected embryos and a stereotypic insertion/deletion (indel) pattern of mutations with the majority of samples exhibiting a 2bp deletion (17). However, we noted that in 5 of the samples analysed, the genotype could not be determined because of failures to PCR amplify the on-target genomic fragment. This finding suggested complexity at the on-target region that may have abolished one or both PCR primer binding sites. Moreover, we identified that 57 of the 137 successfully genotyped samples (42%) exhibited a homozygous wild-type genotype based on PCR amplification of a short genomic fragment (17). We originally interpreted these cases as unsuccessful targeting events, however, given the frequencies of the on-target complexities noted above, we speculated that our previous methods may have missed more complex on-target events.

Here, we have developed computational pipelines to analyse single-cell low-pass whole genome sequencing (WGS), transcriptome and deep-amplicon sequencing data to assess the prevalence of LOH events in the context of CRISPR-Cas9-edited early human embryos (Fig. S1). Our results indicate that LOH events on chromosome 6, including chromosomal and segmental copy number abnormalities, are more prevalent in OCT4-edited embryos compared to both Cas9-injected and uninjected controls, adding to the growing body of literature reporting that CRISPR-Cas9 genome editing can cause unintended on-target damage. Altogether, this underscores the importance of evaluating genome-edited samples for a diversity of mutations, including large-scale deletions, complex rearrangements and cytogenetic abnormalities, undetectable with methods that have routinely been used to interrogate targeted sites in previous studies. Our results sound a note of caution for the potential use of the CRISPR-Cas9 genome editing technology described here for reproductive purposes.

## Results

### Segmental losses and gains at a CRISPR-Cas9 on-target site identified by cytogenetics analysis

In our previous study (17), *in vitro* fertilised zygotes donated as surplus to infertility treatment were microinjected with either an sgRNA-Cas9 ribonucleoprotein complex to target *POU5F1* or Cas9 protein alone as a control and cultured for up to 6 days (targeted and control samples, respectively). We collected a single cell or a cluster of 2-5 cells from these embryos for cytogenetic, genotyping or transcriptomic analysis (Fig. S1).

To determine whether CRISPR-Cas9 genome editing leads to complex on-target DNA damage that would have been missed by our previous targeted amplicon sequencing, we reanalysed low-pass WGS data following whole-genome amplification (WGA) from 23 OCT4-targeted and 8 Cas9 control samples (*SI Appendix,* Table S1). Given the small sample size, we microinjected additional human embryos with a ribonucleoprotein complex to target *POU5F1*, or the Cas9 enzyme as a control, followed by single-cell WGA and low-pass WGS, as before (17). Here and below, the prefix that distinguishes the processing steps is followed by an embryo number and a cell number. The samples used for low-pass WGS were identified with prefix L_ (Fig. S1). The letter C precedes the embryo number to distinguish CRISPR-Cas9 targeted from control samples (Fig. S1). Low-pass WGS data were used to generate copy number profiles for each sample to investigate the presence of abnormalities with a focus on chromosome 6 (Fig. 1A). As an additional comparison, we performed single-cell WGA and low-pass WGS of uninjected control embryos and distinguish these samples with a letter U preceding the embryo number (Fig. S1)

**Fig. 1.**
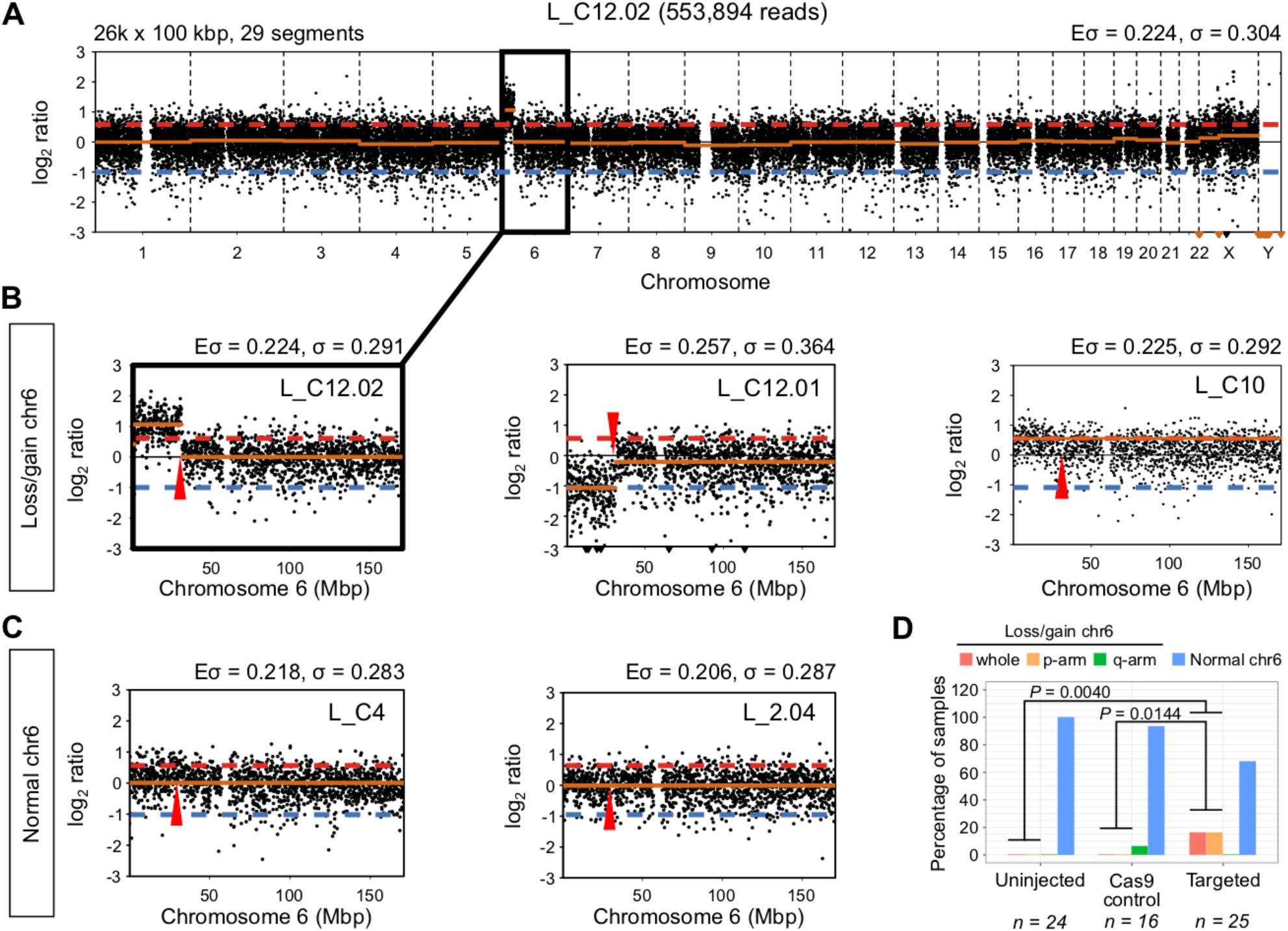
Segmental losses/gains of chromosome 6 are prevalent in OCT4-targeted embryo samples. *(A)* Copy number profile of sample L_C12.02. The segmental gain of chromosome 6 is highlighted. The profile was constructed with 26,000 bins of size 100 kbp, which produced 29 segments. The expected (Eσ) and measured (σ) standard deviation of the profile are reported. *(B)* Zoomed-in view of the copy number profile for samples with segmental losses or gains of chromosome 6. *(C)* Zoomed-in view of the copy number profile for samples with normal chromosome 6. The Eσ and σ reported in B and C correspond to the chromosome only. The approximate position of the *POU5F1* gene is indicated by a red arrow. The red dashed line indicates a copy ratio of 3:2, while the blue dashed lines corresponds to a copy ratio of 1:2. *(D)* The percentage of control and targeted samples with whole or segmental losses/gains of chromosome 6 according to their copy number profiles. P-values are the result of two-tailed Fisher’s tests.

After pre-processing and quality control, we examined the profiles of 65 samples (25 CRISPR-Cas9 targeted, 16 Cas9 controls, and 24 uninjected controls Figs. S2A and S2B). 56 samples exhibited two copies of chromosome 6 with no obvious cytogenetic abnormalities (Figs. 1C, 1D and S3-S5). 17 of the CRISPR-Cas9 targeted samples, or 68%, had no evidence of abnormalities on chromosome 6. By contrast, we observed that 8 out of the 25 targeted samples had evidence of abnormalities on chromosome 6. 4 targeted samples presented a segmental loss or gain that was directly adjacent to or within the *POU5F1* locus on the p-arm of chromosome 6 (Figs. 1B, 1D and S5). Interestingly, this included two cells from the same embryo where one exhibited a segmental gain and the other a reciprocal loss extending from 6p21.3 to the end of 6p (Fig. 1B). Altogether, segmental abnormalities were detected in 16% of the total number of CRISPR-Cas9 targeted samples that were evaluated. We also observed that 4 targeted samples had evidence of a whole gain of chromosome 6 (Figs. 1B, 1D and S5), which also represents 16% of the targeted samples examined. Conversely, a single Cas9 control sample (6.25%) had evidence of a segmental gain on the q-arm of chromosome 6, which was at a site distinct from the *POU5F1* locus (Fig. S4). The uninjected controls did not display any chromosomal abnormalities (Figs. 1D and S3).

The number of segmental and whole-chromosome abnormalities observed in the CRISPR-Cas9 targeted human cells was significantly different from that in the Cas9 (*P* = 0.0144, two-tailed Fisher’s test) and uninjected control (*P* = 0.0040, two-tailed Fisher’s test) samples (Fig. 1D). Moreover, this significant difference can be attributed to the observed segmental abnormalities on 6p, because excluding them from the comparison results in a negligible difference in whole-chromosome abnormalities between targeted and Cas9 control samples (*P* = 0.1429, two-tailed Fisher’s test). This conclusion is further supported by the fact that none of the targeted samples show segmental losses or gains on the p-arm of chromosomes 5 and 7, the closest in overall size to chromosome 6, but the frequency of whole chromosome abnormalities is similar to that observed for chromosome 6, suggesting that genome editing does not exacerbate the rates of whole chromosome errors (Fig. S2C). The comparison we performed between Cas9 control and CRISPR-Cas9 genome edited samples includes a combination of both cleavage and blastocyst stage samples (Table S1). Because rates of aneuploidy are known to be significantly higher at the cleavage stage compared to the blastocyst (18), we wondered whether excluding the samples at the earlier cleavage stage would alter the conclusions drawn about the rates of aneuploidy in CRISPR-Cas9 targeted cells. Here, we found that in comparison to uninjected controls there remained a significantly higher proportion of chromosome 6 aneuploidies in OCT4-targeted cells collected at the blastocyst stage (Fig. S2D). Altogether, low-pass WGS analysis suggests that a significant proportion of unexpected on-target events leads to segmental abnormalities following CRISPR-Cas9 genome editing in human preimplantation embryos.

### Loss-of-heterozygosity identified by targeted deep sequencing

The copy-number profiles described above with low-pass WGS data can only provide a coarse-grained karyotype analysis. To independently investigate the prevalence of LOH events at finer resolution and increased sequencing depth, we designed PCR primer pairs to amplify 15 fragments spanning a ~20kb region containing the *POU5F1* locus. We also included a control PCR amplification in the *ARGFX* locus located on chromosome 3 (*SI Appendix,* Table S4). The PCR amplicons were used to perform deep sequencing by Illumina MiSeq using the gDNA isolated and amplified from 137 single cells or a cluster of 2-5 microdissected cells (111 CRISPR-Cas9 targeted and 26 Cas9 controls) (Fig. S1 and *SI Appendix,* Table S2). The prefix W_ distinguished samples whose gDNA was isolated solely for WGA and the prefix G_ was used to demarcate samples that underwent WGA via the G&T-seq protocol (19). All of these samples were different from the samples used for the cytogenetic analyses above.

We then took advantage of the high coverage obtained at each of the sequenced fragments to call single nucleotide polymorphisms (SNPs), which allowed us to identify samples with putative LOH events: cases in which heterozygous variants, indicative of contribution from both parental alleles, cannot be confidently called in the amplicons flanking the CRISPR-Cas9 on-target site directly. Since we do not have the parental genotype from any of the samples that we analysed, we cannot exclude the possibility that they inherited a homozygous genotype. Therefore, we required the presence of heterozygous SNPs in at least one additional cell from the same embryo to call putative LOH events.

The variant-calling pipeline that we implemented was specifically adjusted for MiSeq data from single cell amplified DNA and includes stringent pre-processing and filtering of the MiSeq reads (Methods). To have sufficient depth of coverage and to construct reliable SNP profiles, we only considered samples with ≥ 5x coverage in at least two thirds of the amplicons across the *POU5F1* locus (Methods and Fig. S6A). This threshold allowed us to retain as many samples as possible and still be confident in SNP calling (20). In addition, we implemented a step in our SNP calling pipeline to control for allele overamplification bias, which is a common issue with single cell amplified DNA (21). This step changes homozygous calls to heterozygous if the fraction of reads supporting the reference allele is above the median value across samples (Figs. S6B and S6C and Methods). Thus, we proceeded with 42 CRISPR-Cas9 targeted and 10 Cas9 control samples with reliable SNP profiles for subsequent analysis. These data led to the identification of four different patterns: samples without clear evidence of LOH, samples with LOH at the on-target site, bookended and open-ended LOH events (Fig. 2A and Figs. S7-S12).

**Fig. 2.**
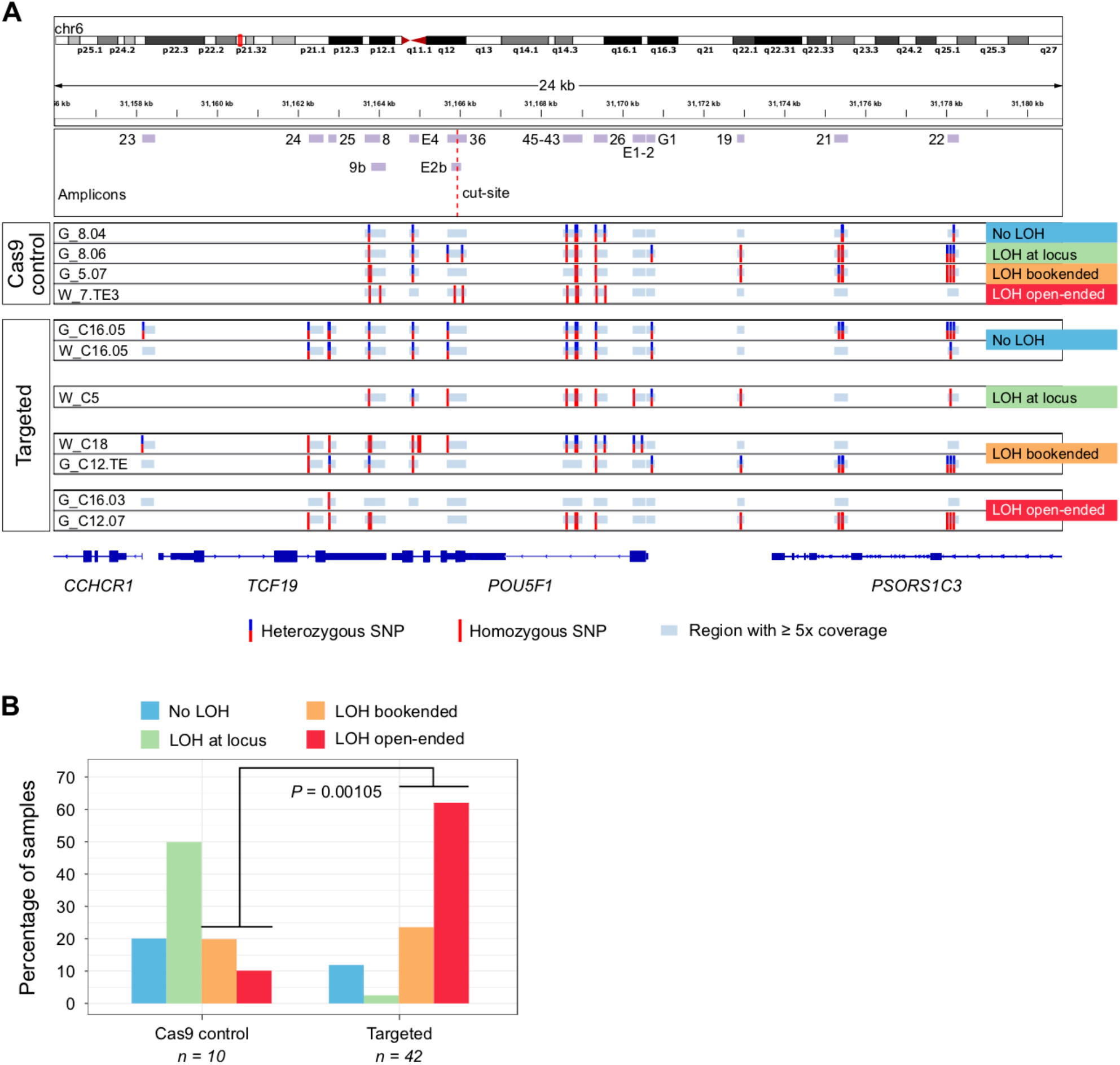
LOH in the *POU5F1* locus is prevalent among OCT4-targeted embryo samples. *(A)* Single nucleotide polymorphism (SNP) profiles constructed from deep sequencing of the depicted amplicons. The four types of loss-of-heterozygosity (LOH) events observed are exemplified. Note that there are amplicons with ≥5x coverage in which SNPs were not called because all reads agree with the reference genome. *(B)* The frequency of each type of LOH event in control and targeted samples. P-value is the result of a two-tailed Fisher’s test.

In samples without LOH (20% of control and 11.9% of targeted samples), we were able to call heterozygous SNPs in multiple amplified fragments (G_8.04, G_C16.05 and W_C16.05, Fig. 2A). Cases with putative LOH at the locus have heterozygous SNPs in the amplicons covering exons 1 and 5 of the *POU5F1* gene (fragments E1-2, G1 and E4 in Fig. 2A) and homozygous SNPs in between (50% of control and 2.4% of targeted samples). These putative LOH samples would have had to have a cell isolated from the same embryo that had a detectable SNP(s) anywhere in between these flanking exons (e.g. see samples G_8.03 versus G_8.04 in Fig. S7). Interestingly, this was the most prevalent pattern in Cas9 control samples (Fig. 2B and Fig. S7), which may indicate the possibility of technical issues due to sequencing or overamplification of one parental allele (see below). Bookended samples have two heterozygous SNPs flanking the cut site but in fragments outside the *POU5F1* locus (20% of control and 23.8% of targeted samples). These LOH events could represent deletions of lengths between ~7kb (G_C12.03, Fig. S10) and ~12kb (W_C11.04, Fig. S9). Finally, in open-ended samples (10% of control and 61.9% of targeted samples) it was not possible to find heterozygous SNPs in any of the amplified fragments (G_C12.07, Fig. 2A) or there was one or a few heterozygous SNPs on only one side of the region of interest (G_C16.02, Fig. S12). This was the most common pattern in targeted samples (Fig. 2B and Figs. S8-S12) and could represent large deletions of ~20kb in length (the size of the region explored) or larger.

As mentioned above, the MiSeq data must be interpreted with caution given the presence of “LOH events” in Cas9 controls. The gDNA employed in these experiments was extracted and amplified with a kit based on multiple displacement amplification (MDA, Methods), which is common in single cell applications but is known to have high allelic dropout and preferential amplification rates (22). Even though, as mentioned above, we implemented a step to control for these biases, this estimate likely under-calls samples with heterozygosity. For example, some homozygous SNPs had 5% of reads mapping to the reference allele but remained homozygous because they fall below the threshold that we used. Considering that we lack the parental genotypes as a reference to choose a more informed cut-off, our method to calculate one from the data represents an unbiased means to correct the presumed allele over-amplification in the samples. Moreover, we cannot exclude the possibility that the analysed single cells inherited a homozygous genotype in the explored region. Nevertheless, the fact that there is a significant number of CRISPR-Cas9 targeted samples with the largest LOH patterns is notable (Fig. 2B).

### Unexpected CRISPR-Cas9-induced on-target events do not lead to preferential misexpression of telomeric to POU5F1 genes

Our low-pass WGS and SNP analysis above indicate mutations at the *POU5F1* locus that are larger than discrete indels. We therefore wondered if this on-target complexity may encompass the mutations of genes adjacent or telomeric to *POU5F1* that could complicate the use of CRISPR-Cas9 to understand gene function in human development or other contexts where the analysis of primary cells is required. To address this, we reanalysed the single-cell RNA sequencing (scRNA-seq) transcriptome datasets (Table S6) we generated previously (17) and focused on the chromosome location of transcripts (Figs. 3A-C). This analysis indicated that differentially expressed genes are not biased to a specific chromosome (Fig. 3A). Moreover, differentially expressed genes are not enriched to either chromosome 6 or the region telomeric to the CRISPR-Cas9 on-target site (Fig. 3D). These results suggest that the transcriptional differences observed as a consequence of *POU5F1* targeting are not confounded by mutations of genes adjacent, or telomeric, to the on-target locus. This could be due to a number of reasons. For example, given that the proportion of samples that exhibit unintended CRISPR-Cas9-induced mutations (e.g. segmental aneuploidies or LOH events) is low, the sample size used is sufficiently high to mask any transcriptional differences in genes adjacent to the cut site in samples with segmental loss of the p-arm of chromosome 6. It is also possible that the extent of the on-target complexity is exaggerated using the gDNA-based pipelines we developed. Notably, because we use single-cell samples, as mentioned above, these are prone to allele over-amplification and this can confound the interpretation of on-target mutation complexity.

**Fig. 3.**
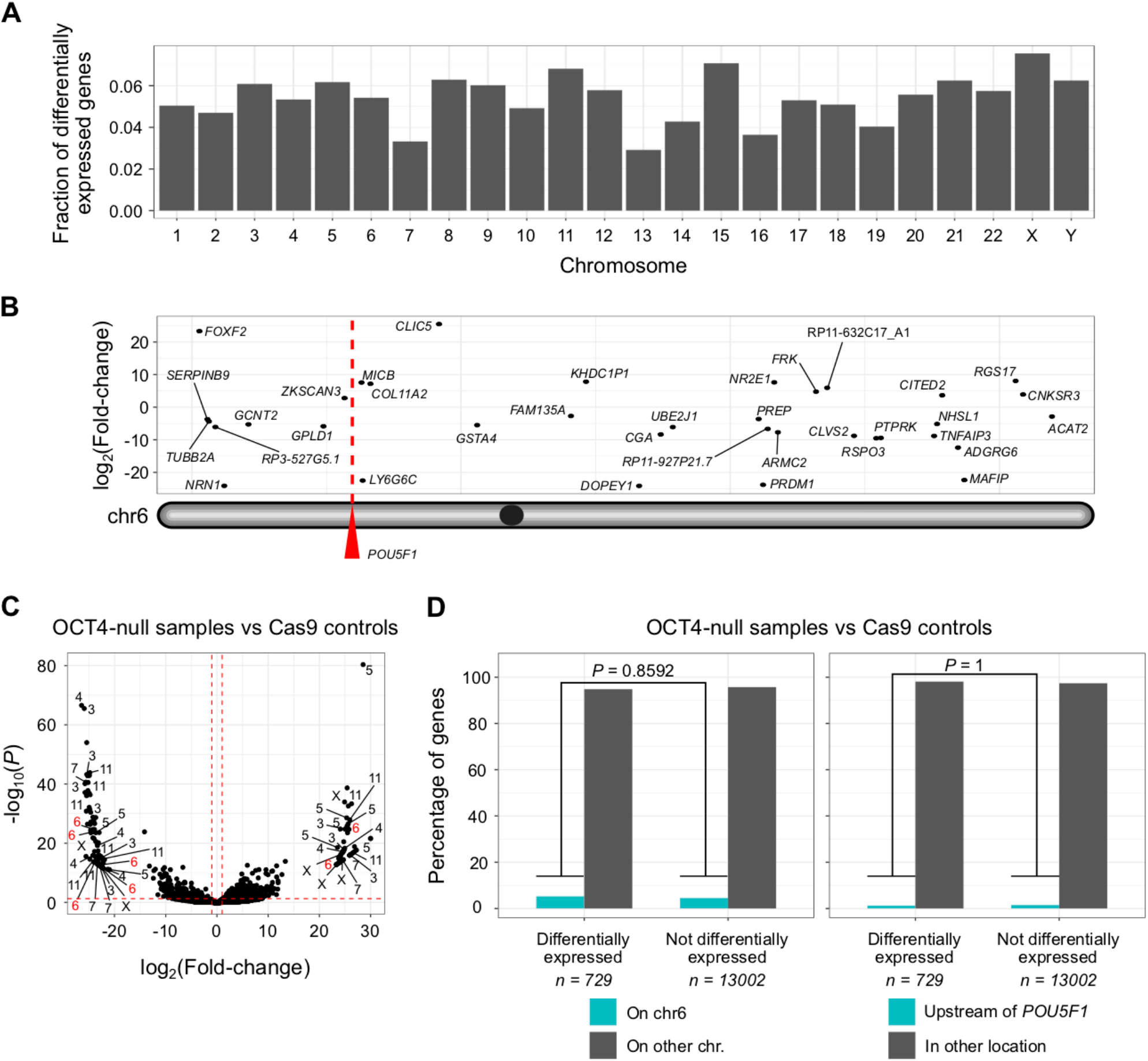
LOH in OCT4-targeted samples does not lead to preferential misexpression of genes located on chromosome 6. *(A)* The fraction of differentially expressed genes per chromosome from the comparison between OCT4-null samples and Cas9 controls. *(B)* Location of differentially expressed genes along chromosome 6. *(C)* Volcano plot summarising the comparison between OCT4-null samples and Cas9 controls with differential gene expression analysis. The chromosome location of some of the most dysregulated genes is shown (absolute log2 fold change > 20 and Benjamini-Hochberg adjusted *P* < 0.05). The red dashed lines correspond to absolute log2 fold changes > 1 and Benjamini-Hochberg adjusted *P* < 0.05. *(D)* Genes located on chromosome 6 are not overrepresented in the list of loci whose expression is disturbed upon OCT4 knock out. The same applies for genes directly upstream to the *POU5F1* gene. P-values are the result of two-tailed Fisher’s tests.

### No evidence of on-target complexity using digital karyotype and LOH analysis of the single-cell transcriptome data

The use of RNA-seq data to detect chromosomal abnormalities (23) has great potential to complement the informative low-pass WGS or array CGH methods currently used for embryo screening in the context of assisted reproductive technologies (24, 25). In addition to karyotype analysis, transcriptome data may also provide information about embryo competence at the molecular level. Groff and colleagues have demonstrated that aneuploidy can be estimated based on significant variations in gene expression in the affected chromosome(s) compared to reference control samples (24). In addition, Weissbein *et al.* developed a pipeline, called eSNP-Karyotyping, for the detection of LOH in chromosome arms (26). eSNP-Karyotyping is based on measuring the ratio of expressed heterozygous to homozygous SNPs. We applied these two approaches, hereinafter referred to as z-score- and eSNP-Karyotyping, to the single-cell RNA-seq (scRNA-seq) samples (distinguished with the prefix T_) obtained using the G&T-seq protocol (14) (*SI Appendix*, Table S3). This allowed us to investigate whether transcriptome data could be used to determine the frequency of LOH events in CRISPR-Cas9 targeted embryos.

Since eSNP-Karyotyping relies on SNP calls from gene expression data, it is very sensitive to depth and breadth of sequencing (26). Therefore, we used results from this method as a reference to select high quality samples for our transcriptome-based analyses (Fig. S13A-C). After these filtering steps, we retained 38 samples (22 CRISPR-Cas9 targeted and 16 Cas9 controls) to analyse further.

In general, we found good agreement between the chromosomal losses detected by z-score-karyotyping and the LOH events identified by eSNP-Karyotyping (Fig. S14A and S14B). For example, the digital karyotype of Fig. S14A shows the loss of chromosome 4, the p-arm of chromosome 7 and the q-arm of chromosome 14 in sample T_7.01, as well as the loss of chromosome 3 and the p-arm of chromosome 16 in sample T_C16.06. These abnormalities are identified as LOH events in the eSNP-Karyotyping profiles of the same samples (Fig. 14B). Moreover, the copy number profiles built from low-pass WGS data for different cells from the same embryos also corroborates these chromosomal abnormalities (Fig. S13D and S13E). In terms of events that could be associated with CRISPR-Cas9 on-target damage, z-score-karyotyping identified the loss of chromosome 6 in sample T_C12.07 (Fig. 4A), which is consistent with the open-ended LOH pattern observed in the gDNA extracted from the same cell G_C12.07 (Fig. S10) and the segmental loss detected in sample L_C12.01 from the same embryo (Fig. 1B). Also, the gain of the p-arm of chromosome 6 was detected in sample T_C12.15 (Fig. 4A), which is consistent with the segmental gain observed in sample L_C12.02 from the same embryo (Fig. 1B). The gains and losses of chromosome 6 in samples T_2.02, T_2.03, T_2.14, T_7.02 and T_C16.06 (Fig. 4A) are difficult to interpret due to the low quality of their MiSeq data or the lack of amplicon information for the q-arm (Fig. S7 and Fig. S12). Interestingly, eSNP-Karyotyping did not detect any LOH events in chromosome 6 (Fig. S15), suggesting that this approach is not sensitive enough to detect segmental abnormalities in single cell samples. Overall, the transcriptome-based karyotypes did not confirm the trends observed in the gDNA-derived data (Fig. 4B).

**Fig. 4.**
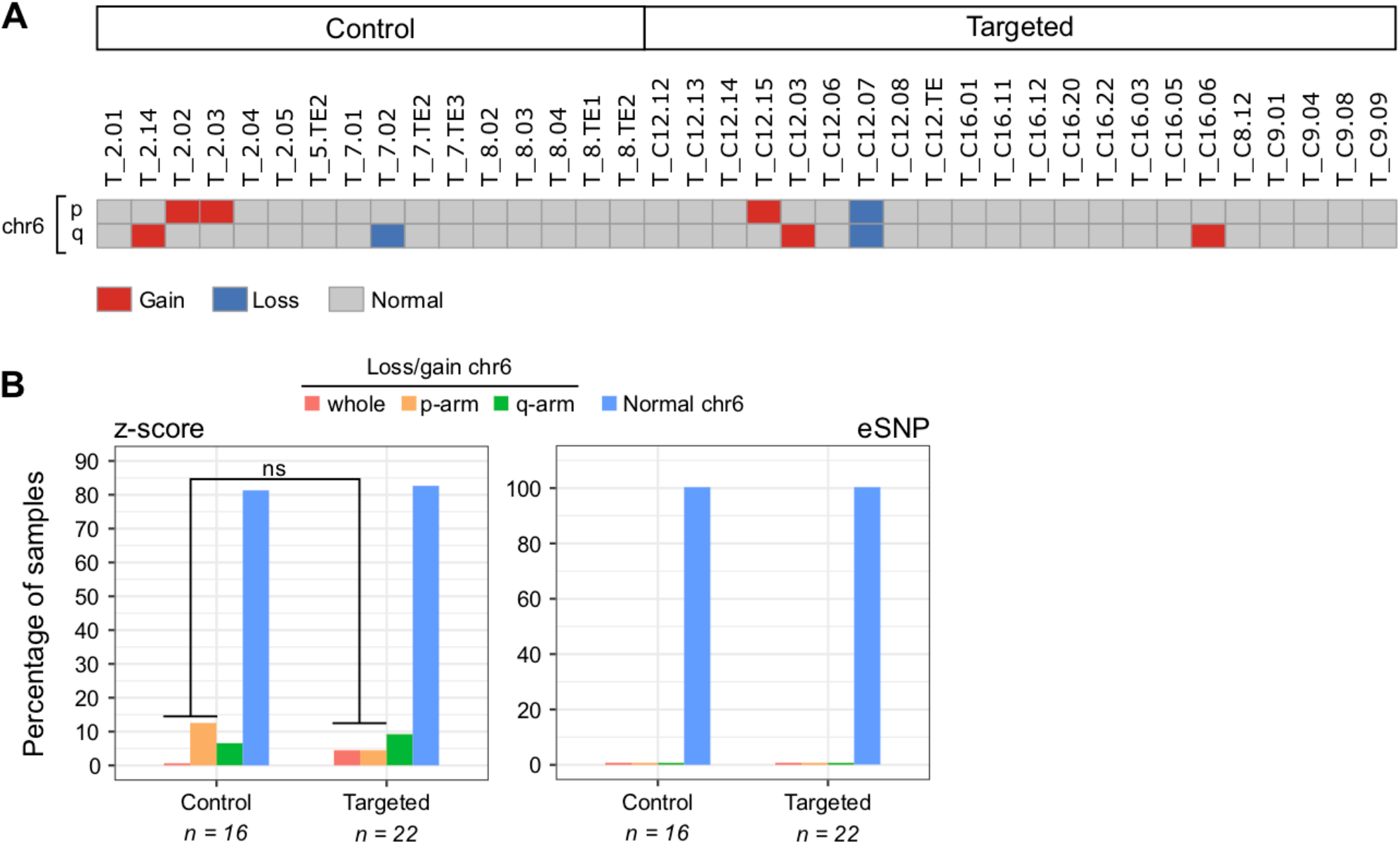
Transcriptome-based karyotypes do not capture segmental losses/gains of chromosome 6 in OCT4-targeted embryo samples. *(A)* Digital karyotype based on the total gene expression deviation from the average of each chromosome arm (z-score-karyotyping). Only chromosome 6 (see Fig. S14A for the rest of the chromosomes). *(B)* The percentage of control and targeted samples with segmental losses/gains of chromosome 6 according to their transcriptome-based karyotype (see Figs. S14A and S15). P-value is the result of a two-tailed Fisher’s test.

## Discussion

In all, we reveal unexpected on-target complexity following CRISPR-Cas9 genome editing of human embryos. Our data suggest approximately 16% of samples exhibit segmental losses/gains adjacent to the *POU5F1* locus and LOH events that span 4kb to at least 20kb. Chromosome instability, including whole or segmental chromosome gain or loss, is common in human preimplantation embryos (27, 28). However, in contrast to Cas9 control embryos, we noted a significantly higher frequency of CRISPR-Cas9 targeted embryos with a segmental gain or loss that was directly adjacent to the *POU5F1* on-target site. The segmental errors were observed in embryos from distinct genetic backgrounds and donors. Therefore, together with their on-target location, this suggests that the errors may have been an unintended consequence of CRISPR-Cas9 genome editing. This is supported by the higher frequency of larger LOH events that we observed in CRISPR-Cas9 targeted embryos compared to Cas9 controls using an independent targeted deep-sequencing approach. However, due to the nature of our datasets (shallow sequencing, MDA-amplified gDNA, lack of parental genotypes) we may be overestimating LOH events. This may explain some of the on-target complexity observed in Cas9 control samples but does not account for the significantly higher proportion of LOH in the CRISPR-Cas9 targeted samples. It is important to note that 68% of CRISPR-Cas9 targeted cells did not exhibit any obvious segmental or whole chromosome 6 abnormalities, indicating that their genotype and phenotype, with respect to OCT4 function, are interpretable. Moreover, our transcriptome-based digital karyotypes and differential gene expression analysis indicate biallelic transcripts and gene expression up- and down-stream of the *POU5F1* locus in so far as is resolvable from scRNA-seq data, suggesting that in these samples the LOH does not lead to the misexpression of other genes adjacent to the *POU5F1* locus. Also, our work and previous accounts of unexpected CRISPR-Cas9 editing outcomes (9, 10, 12–14, 16) indicate that the frequency of discrete on-target events predominates, which should increase the confidence of the interpretation of functional studies in human embryos. Given the likelihood of mosaicism, it is unclear whether the segmental abnormalities we observed in any one cell analysed from each embryo are representative of the entire CRISPR-Cas9 targeted embryo or a subset of cells within the embryo. Altogether, this points to the need to use robust techniques to distinguish cells affected by on-target complexity and large deletions following CRISPR-Cas9-mediated genome editing from cells with less complex mutations and our computational pipelines and multi-omics analyses are approaches that may be used in the future.

By contrast, we did not observe significantly more abnormalities on chromosome 6 using methods to determine LOH or karyotype from scRNA-seq datasets. There are several factors that could account for the discrepancy between these datasets. Firstly, we do not have the transcriptome from the same samples that showed gains and losses of chromosome 6 in the cytogenetics analysis. A follow-up study in which both transcriptomics and cytogenetics data are extracted from the same sample would be very informative and could be performed by modifying the G&T-seq protocol (19) to incorporate a multiple annealing and looping-based amplification cycles (MALBAC) method for WGA (29) in place of MDA, which was used here due to the proofreading activity of the phi29 MDA polymerase at the expense of high preferential amplification rates (22). Secondly, mosaicism is common in human preimplantation embryos (30) and this could explain why the digital karyotypes based on gene expression did not detect abnormalities at the same rate as the copy number profiles. Another possibility is that the LOH events are not sufficiently large to impact total gene expression of chromosome 6, which is what z-score- and eSNP-Karyotyping rely on. This could also account for the cytogenetics results, as LOH up to a few Mb in size could cause mapping issues due to the very low coverage of shallow sequencing that are reflected as gains and losses of whole chromosome segments. Finally, the LOH events detected by gDNA-derived data may only affect genes that are not expressed in the embryo context or whose expression is so low that it cannot be accurately measured by scRNA-seq. So, when z-score- and eSNP-Karyotyping compare gene or SNP expression of targeted versus control samples, no significant differences are identified.

The segmental aneuploidies identified by cytogenetics analysis (Figs. 1B and S3-S5) most probably point to the occurrence of complex genomic rearrangements in OCT4-targeted samples, such as chromosomal translocations or end-to-end fusions, as it seems unlikely that the rest of the chromosome would continue to be retained without a telomere (31–33). It is likely that human embryos tolerate aneuploidy up to embryo genome activation, given that even embryos with observed multipolar spindles continue to develop during early cleavage divisions (34). Following this, chromosomal anomalies are likely to become increasingly detrimental to cellular viability, although a degree of tolerance may persist in trophectoderm cells (28). Why early embryos fail to arrest despite chaotic chromosomal errors such as multipolar spindle formation or presumptive unresolved double strand breaks following CRISPR-Cas9 genome editing is unclear and crucial to understand. An important next step to gain insights into the extent of the damage would be to use alternative methods. One possibility to understand the complexity would be to perform cytogenetic analysis using fluorescence *in situ* hybridization (FISH) (35) to probe for segments of chromosome 6. Another option is a chromosome walk-along approach to amplify genomic fragments even further away from the 20kb genomic region that we evaluated, in order to bookend heterozygous SNPs on either side of the *POU5F1* on-target site. This may be kilo- or mega-bases away from the on-target site based on previous publications in the mouse or human cell lines (9, 10, 12–14).

Based on our data, the possibility of gene editing via IH-HR cannot be definitely excluded. A pre-print by Liang *et al.* (36) suggests that IH-HR could be one of the major DNA double-strand break repair pathways in human embryos. Following a similar approach to their previous study (8), the authors used CRISPR-Cas9-mediated genome editing to target a paternal mutation and were able to amplify an ~8kb genomic DNA fragment which, together with G-banding and FISH of ESCs derived from targeted embryos, suggests that repair from the maternal chromosome by IH-HR results in a stretch of LOH. Of note, due to the selection bias that occurs during ESC derivation and the mosaicism observed following genome editing, it is not possible to draw definitive conclusions about the extent of LOH or its cause in an embryo context, whereby cells with complex mutations may be preferentially excluded from ESC derivation. By contrast, another pre-print by Zuccaro *et al.* using the same microinjection method suggests that the LOH observed following CRISPR-Cas9-mediated genome editing is a consequence of whole chromosome or segmental loss adjacent to the on-target site and that microhomology-mediated end-joining (MMEJ) is the dominant repair pathway in this context (37). This corroborates our previous findings in human embryos targeted post-fertilisation, where we noted a stereotypic pattern to the type of indel mutations and speculated that this was likely due MMEJ (17). Although microhomologies can promote gene conversion by, for example, inter-chromosomal template switching in a RAD51-dependent manner (38), based on our previous transcriptome analysis, we found that components of the MMEJ pathway (i.e. *POLQ*) are transcribed in early human embryos, while factors essential for HDR (i.e. *RAD51*) are not appreciably expressed. This suggests that MMEJ-derived large deletions (14, 37) are more likely than microhomology-mediated gene conversion in this context, though protein expression has yet to be fully characterised. Consistent with this, a significant fraction of somatic structural variants arises from MMEJ in human cancer (39). Moreover, microhomology-mediated break-induced replication underlies copy number variation in mammalian cells (40) and microhomology/microsatellite-induced replication leads to segmental anomalies in budding yeast (41). The discrepancy between the Liang *et al.* and Zuccaro *et al.* studies could be due to locus-dependent differences of CRISPR-Cas9 genome editing fidelity. For example, Przewrocka *et al.* demonstrate that the proximity of the CRISPR-Cas9-targeted locus to the telomere significantly increases the possibility of inadvertent chromosome arm truncation (16). To fully elucidate the LOH that has occurred at the on-target site in our study, and to resolve the controversy over the IH-HR reported by others (8, 9, 36, 37), will require the development of a pipeline to enrich for the region of interest and then perform deep (long-read) sequencing to evaluate the presence and extent of on-target damage. By bookending SNPs on either side of an LOH event, primers could be designed to incorporate the SNPs and ensure that both parental alleles are amplified. However, this is difficult to perform, and alternative methods include using CRISPR gRNAs to cut just outside of the LOH region followed by long-read sequencing (42).

It would also be of interest to evaluate whether other genome editing strategies, such as prime and base editing, nickases or improvements in the efficiency of integrating a repair template, may reduce the on-target complexities observed by us and others using spCas9. However, non-negligible frequencies of editing-associated large deletions have been reported after the use of the Cas9^D10A^ nickase in mESCs (14) and prime editing in early mouse embryos (43). By contrast, while proof-of-principle studies suggest that base editors could be used to repair disease-associated mutations in human embryos, further refinements to reduce the likelihood of unexpected conversion patterns and high rates of off-target edits would be of benefit (2). There are too few studies to date using repair templates. Of the studies that have been conducted, the reported efficiencies of repair with templates in human embryos are very low (5, 7, 8). Modulation of DNA damage repair factors or tethering Cas9 enzymes with a repair template may yield improvements that could allow for the control of editing outcomes.

Our re-evaluation of on-target mutations, together with previous accounts of unexpected CRISPR-Cas9 on-target damage (9, 10, 12–14), strongly underscores the importance of further basic research in a number of cellular contexts to resolve the damage that occurs following genome editing. Moreover, this stresses the significance of ensuring whether one or both parental chromosome copies are represented when determining the genotype of any sample to understand the complexity of on-target CRISPR mutations, especially in human primary cells.

## Methods

### Ethics statement

We reprocessed the DNA and reanalysed the data generated in our previous study (17). This corresponds to 168 samples (134 OCT4-targeted and 34 Cas9 controls) across 32 early human embryos (24 OCT4-targeted and 8 Cas9 controls). For the present work, we used 56 additional single-cell samples (19 OCT4-targeted, 12 Cas9 controls and 25 uninjected controls) across 22 early human embryos (1 OCT4-targeted, 1 Cas9 control and 20 uninjected controls). This study was approved by the UK Human Fertilisation and Embryology Authority (HFEA): research licence number R0162, and the Health Research Authority’s Research Ethics Committee (Cambridge Central reference number 19/EE/0297). Our research is compliant with the HFEA code of practice and has undergone inspections by the HFEA since the licence was granted. Before giving consent, donors were provided with all of the necessary information about the research project, an opportunity to receive counselling and the conditions that apply within the licence and the HFEA Code of Practice. Specifically, patients signed a consent form authorising the use of their embryos for research including genetic tests and for the results of these studies to be published in scientific journals. No financial inducements were offered for donation. Patient information sheets and the consent documents provided to patients are publicly available (https://www.crick.ac.uk/research/a-z-researchers/researchers-k-o/kathy-niakan/hfea-licence/). Embryos surplus to the IVF treatment of the patient were donated cryopreserved and were transferred to the Francis Crick Institute where they were thawed and used in the research project.

### CRISPR-Cas9 targeting of *POU5F1*

We analysed single cells or trophectoderm biopsies from human preimplantation embryos that were CRISPR-Cas9 edited in our previous study (17) plus an additional 56 samples used in the present work. *In vitro* fertilised zygotes donated as surplus to infertility treatment were microinjected with either a sgRNA-Cas9 ribonucleoprotein complex or with Cas9 protein alone and cultured for 5-6 days (targeted and control samples, respectively). The sgRNA was designed to target exon 2 of the *POU5F1* gene and experiments performed as previously described (17). Genomic DNA from Cas9 control and OCT4-targeted was isolated using the REPLI-g Single Cell Kit (QIAGEN, 150343). DNA samples isolated for cytogenetic analysis were amplified with the SurePlex Kit (Rubicon Genomics). See the *SI Appendix* for more details.

### Cytogenetic analysis

Low-pass whole genome sequencing (depth of sequencing < 0.1x) libraries were prepared using the VeriSeq PGS Kit (Illumina) or the NEB Ultra II FS Kit and sequenced with the MiSeq platform as previously described (17) or with Illumina HiSeq 4000, respectively. Reads were aligned to the human genome hg19 using BWA v0.7.17 (44) and the copy number profiles generated with QDNAseq v1.24.0 (45). See the *SI Appendix* for more details.

### PCR primer design and testing

PCR primer pairs were designed with the Primer3 webtool (http://bioinfo.ut.ee/primer3/, Table S4). We restricted the product size to 150-500bp and used the following primer temperature settings: Min=56, Opt=58, Max=60. We tested all primers using 1uL of genomic DNA from H9 human ES cells in a PCR reaction containing 12.5 uL Phusion High Fidelity PCR Master Mix (NEB, M0531L), 1.25 uL 5 uM forward primer, 1.25 uL 5 uM reverse primer and 9 uL nuclease-free water. Thermocycling settings were: 95°C 5min, 35 cycles of 95°C 30s, 58°C 30s, 72°C 1min, and a final extension of 72°C 5min. We confirmed the size of the PCR products by gel electrophoresis. See the *SI Appendix* for more details.

### PCR amplification and targeted deep sequencing

Isolated DNA was diluted 1:100 in nuclease-free water. We used the QIAgility robot (QIAGEN, 9001531) for master mix preparation (see above) and distribution to 96-well plates (Table S5). Then, the Biomek FX liquid handling robot (Beckman Coulter, 717013) was used to transfer 1uL of DNA to the master mix plates and to mix the reagents. The PCR reaction was run with the settings described above. PCR products were cleaned with the Biomek FX robot using the chemagic SEQ Pure20 Kit (PerkinElmer, CMG-458). Clean PCR amplicons from the same DNA sample were pooled to generate 137 libraries that were sequenced by Illumina MiSeq v3. See the *SI Appendix* for more details.

### SNP-typing

We trimmed the MiSeq paired-end reads with DADA2 (46), corrected substitution errors in the trimmed reads with RACER (47) and mapped the corrected reads to the human genome hg38 with BWA v0.7.17 (44). Subsequently, SAM files were converted to the BAM format and post-processed using Samtools v1.3.1 (48). SNP calling was performed with BCFtools v1.8 (49) using mpileup and call. SNPs supported by less than 10 reads and with mapping quality below 50 were filtered out. To control for allele overamplification, homozygous SNPs were changed to heterozygous if the fraction of reads supporting the reference allele was at least 6% of the total (21). This threshold corresponds to the median of the distribution of the fraction of reads supporting the reference allele across samples. See the *SI Appendix* for more details.

### scRNA-seq data analysis

scRNA-seq reads from G&T-seq samples were processed as previously described (17). Samples with a breadth of sequencing below 0.05 were not considered for any downstream analysis (Fig. S13A-C). Differential gene expression analysis was carried out with DESeq2 v1.10.1 (50). For digital karyotyping based on gene expression, we adapted the method described in (24) to identify gains or losses of chromosomal arms (z-score-karyotyping). For digital karyotyping based on SNP expression, we applied the eSNP-Karyotyping pipeline with default parameters (26). See the *SI Appendix* for more details.

### Data and software availability

All data supporting the findings of this study are available within the article and its supplementary information. MiSeq and low-pass WGS data have been deposited to the Sequence Read Archive (SRA) under accession number PRJNA637030. scRNA-seq data was extracted from the Gene Expression Omnibus (GE) using accession GSE100118. A detailed analysis pipeline is available at the following site: https://github.com/galanisl/loh_scripts.

## Acknowledgements

We thank the generous donors whose contributions have enabled this research. We thank Robin Lovell-Badge, James Haber, Alexander Frankell, Aska Przewrocka, Charles Swanton, Maxime Tarabichi, the Niakan and Turner laboratories for discussion, advice and feedback; the Francis Crick Institute’s core facilities including Jerome Nicod and Robert Goldstone at the Advanced Sequencing Facility; D.W. was supported by the National Institute for Health Research (NIHR) Oxford Biomedical Research Centre Programme. N.K. was supported by the University of Oxford Clarendon Fund and Brasenose College Joint Scholarship. Work in the K.K.N. and J.M.A.T. labs was supported by the Francis Crick Institute, which receives its core funding from Cancer Research UK (FC001120 and FC001193), the UK Medical Research Council (FC001120 and FC001193), and the Wellcome Trust (FC001120 and FC001193). Work in the K.K.N. laboratory was also supported by the Rosa Beddington Fund.

## Author contributions

K.K.N. conceived the project. N.M.E.F. generated the genomics and transcriptomics datasets. A.M., E.H. and G.A-L. designed and tested primers. N.K. and D.W. generated the low-pass WGS data. M.G. performed the PCR amplification experiments with the robotics equipment. J.Z. implemented the variant calling pipeline for the amplicon sequencing data. G.A-L. collected, processed and analysed all the datasets. J.M.A.T. provided advice on the project design. G.A-L. and K.K.N. wrote the manuscript with the help from all other authors. All authors assisted with experimental design and figures.

## Notes

https://www.ncbi.nlm.nih.gov/bioproject/PRJNA637030

### Competing Interest Statement

The authors have declared no competing interest.

