## Supplementary Information for "Frequent loss-of-heterozygosity in CRISPR-Cas9-edited early human embryos"

### Detailed Methods

**CRISPR-Cas9 targeting of *POU5F1*.** The samples that we analysed correspond to single cells or trophectoderm biopsies from human preimplantation embryos that were CRISPR-Cas9 genome edited in our previous study (1) plus an additional 56 samples used in the present work. Briefly, *in vitro* fertilised zygotes that were donated as surplus to infertility treatment were microinjected with either a sgRNA-Cas9 ribonucleoprotein complex or with Cas9 protein alone and cultured for 5-6 days (targeted and control samples, respectively). The sgRNA was designed to target exon 2 of the *POU5F1* gene and experiments performed as previously described (1). Genomic DNA from Cas9 control and OCT4-targeted human embryos was isolated from either an individual single cell or a cluster of 2-5 cells from trophectoderm biopsies from embryos that developed to the blastocyst stage, as well as blastomeres from earlier stage embryos (Table S2 and S3) using the REPLI-g Single Cell Kit (QIAGEN, 150343) according to the manufacturer's guidelines. Since these samples were originally isolated for further processing by G&T-seq (2) or whole genome amplification, they are identified by a G, T or W prefix in Tables S2 and S3. DNA samples isolated for cytogenetic analysis were amplified with the SurePlex Kit (Rubicon Genomics) and are identified by an L prefix (Table S1).

**Cytogenetic analysis.** To determine the chromosome copy number of samples in Table S1, their genomic DNA was subjected to low-pass whole genome sequencing (depth of sequencing < 0.1x). Libraries were prepared using the VeriSeq PGS Kit (Illumina) or the NEB Ultra II FS Kit (Table S1) and sequenced with the MiSeq platform as previously described (1) or with Illumina HiSeq 4000, respectively. Sequenced reads were aligned to the human genome hg19 using BWA version 0.7.17 (3) and the digital karyotypes were generated with the R package QDNAseq version 1.24.0 (4). We used bins of size 100kb and filtered out samples with a strong difference between the measured and expected standard deviations of the generated profile (Fig. S2A and S2B). The expected standard deviation (Eσ) is defined as √1/N, where N is the average number of reads per bin. The measured standard deviation (σ) is calculated from the data with a 0.1%-trimmed first-order estimate (4).

**PCR primer design and testing.** The 15 PCR primer pairs were designed with the Primer3 webtool (http://bioinfo.ut.ee/primer3/) across the *POU5F1* locus (chr6:31,157,800-31,178,600 on hg38, Table S4). We also designed a control primer pair in exon 4 of the gene *ARGFX,* which is on a different chromosome (chr3:121,583,621-121,586,438, Table S4). We restricted the product size to the 150-500bp range and used the following primer temperature settings: Min=56, Opt=58, Max=60. We selected primer pairs with similar melting temperature, length, the lowest possible GC percentage and with amplicons containing at least one common human variation as reported by dbSNP 1.4.2 (https://www.ncbi.nlm.nih.gov/variation/docs/glossary/#common). We tested all primers using 1uL of genomic DNA from H9 human ES cells in a PCR reaction containing 12.5 uL Phusion High Fidelity PCR Master Mix (New England Biolabs, M0531L), 1.25 uL 5 uM forward primer, 1.25 uL 5 uM reverse primer and 9 uL nuclease-free water. Thermocycling settings were: 95°C 5min, 35 cycles of 95°C 30s, 58°C

30s, 72°C 1min, and a final extension of 72°C 5min. We confirmed that the size of the PCR products corresponded to the expected amplicon size (Table S4) by gel electrophoresis.

**PCR amplification.** In preparation for PCR amplification, the DNA isolated from samples in Table S2 was diluted 1:100 in nuclease-free water. To expedite the processing of our 2192 samples (16 target fragments for each of the 137 DNA templates), we used the QIAgility robot (QIAGEN, 9001531) for master mix preparation (see above) and distribution to 96-well plates using the layout depicted in Table S5 for a total of 24 plates. Then, the Biomek FX liquid handling robot (Beckman Coulter, 717013) was used to transfer 1uL of DNA at once to the master mix plates using a 96-multichannel pipetting head and to mix the reagents. The PCR reaction was run with the thermocycling settings described above. PCR products were cleaned with the Biomek FX robot using the chemagic SEQ Pure20 Kit (PerkinElmer, CMG-458) as per manufacturer's instructions.

**Targeted deep sequencing.** Clean PCR amplicons from the same DNA sample were barcoded and pooled to generate 137 barcoded libraries that were submitted for targeted deep sequencing by Illumina MiSeq v3 (300bp paired end reads).

**SNP-typing.** We trimmed the MiSeq paired-end reads with DADA2 (5) to remove low-quality regions (function filterAndTrim with parameters trimLeft=5, truncLen=c(150,150), truncQ=2, maxN=0 and maxEE=c(5, 5)). Then, we corrected substitution errors in the trimmed reads with RACER (6) and mapped the corrected reads to the human genome hg38 with BWA version 0.7.17 (3) in multi-threaded mode using the mem algorithm with default settings. Subsequently, SAM files were converted to the BAM format and post-processed (sorting, indexing and mate fixing) using Samtools version 1.3.1(7). SNP calling was performed with BCFtools version 1.8 (8) using the mpileup (--max-depth 2000 -a 'AD,DP,ADF,ADR' -Ou) and call (-mv -V 'indels' -Ov) algorithms in multi-threaded mode. Since the average length of our amplicons is 300bp and the trimmed reads ended up having length ~145, at least 10 reads are needed to achieve a 5x coverage at each amplicon. Therefore, SNPs supported by less than 10 reads and with mapping quality below 50 were filtered out. To control for allele overamplification, we revisited the homozygous SNP calls in search for reads supporting the reference allele at those positions. We changed these homozygous SNPs to heterozygous if the fraction of reads supporting the reference allele was at least 6% of the total (9). This threshold corresponds to the median of the distribution of the fraction of reads supporting the reference allele across samples. The resulting VCF files were then indexed and inspected in the Integrative Genomics Viewer (10).

**scRNA-seq data analysis.** scRNA-seq reads from G&T-seq samples (Table S3) were aligned to the human reference genome GRCh38 using TopHat2 version 2.1.1 (11). Samples with a breadth of sequencing below 0.05 were not considered for any downstream analysis (Fig. S13A-C). Read counts per gene were calculated using HTSeq 0.12.4 (12) and normalised using TPM units (13). Differential gene expression analysis was carried out with DESeq2 v1.10.1(14). For digital karyotyping based on gene expression, we adapted the method described in (15) to identify gains or losses of chromosomal arms. Briefly, after removal of no-show, mitochondrial, sex chromosome and PAR genes, the TPM expression of all genes mapping to the p-arm of chromosome *i* was summed and compared to the average sum for the same chromosome and arm across samples via the calculation of a z-score. Z-scores with values below -1.65 and above 1.65 were considered segmental losses and gains, respectively. Chromosome arms with values in between were considered to be normal. The same procedure was repeated for the q-arm of each chromosome. For digital karyotyping based on SNP expression, we applied the eSNP-Karyotyping pipeline with default parameters (16). eSNP-Karyotyping identifies loss-of-heterozygosity in a chromosome arm when the ratio of heterozygous to homozygous SNPs in that arm is significantly lower compared to the other chromosome arms. For this, the pipeline employs the GATK best practices for SNP calling using RNA-seq data and compares called heterozygous variants with homozygous variants reported on dbSNP 1.4.2 (16). eSNP-Karyotyping is very sensitive to depth and breadth of sequencing, so we selected samples for our scRNA-seq analyses based on the quality of the eSNP-Karyotyping profiles (Fig. S13A-C and Table S3).

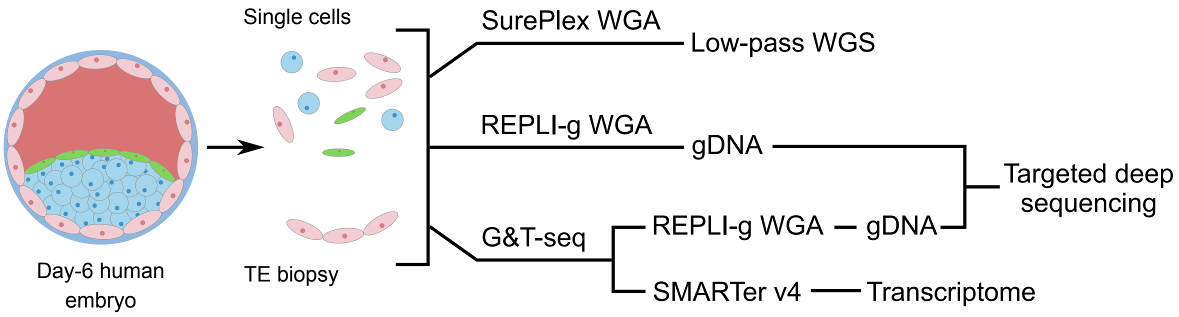


| Targeted cell | Control cell |
|---|---|
| L_C12.01 | L_3.03 (Cas9 control)<br>L_U15 (Uninjected) |
| W_C11.04 | W_7.02 (Cas9 control) |
| G_C16.03 | G_2.14 (Cas9 control) |
| T_C16.03 | T_2.14 (Cas9 control) |

**Fig. S1. Sample types and nomenclature used throughout the paper.** We analysed low-pass whole genome sequencing (WGS) and transcriptome data from OCT4-targeted and Cas9 control single cells or trophectoderm (TE, precursor cells of the placenta) biopsies from human embryo samples. In addition, the genomic DNA (gDNA) isolated from single cells or TE biopsies subjected to the G&T-seq protocol or to whole genome amplification (WGA) was used for targeted deep sequencing across the *POU5F1* locus. Sample identifiers start with a prefix, followed by the embryo and cell number. The embryo number of uninjected samples is preceded by a letter U. The embryo number of CRISPR-edited samples is preceded by a letter C. Prefix L_ corresponds to the low-pass WGS data, prefix W_ to gDNA that was amplified with the REPLI-g kit, prefix G_ to gDNA extracted with the G&T-seq protocol and prefix T_ to scRNA-seq data produced with G&T-seq

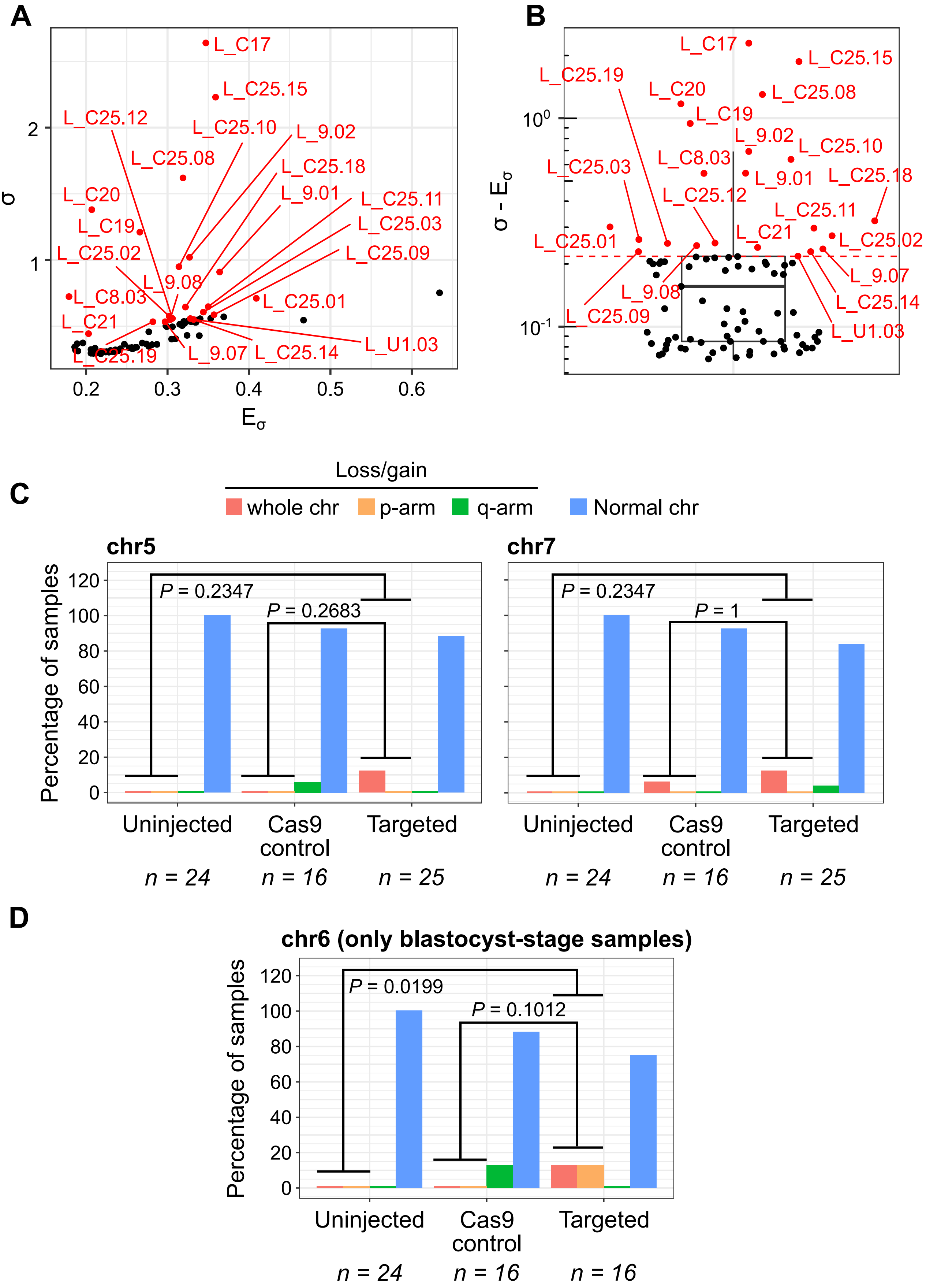


**Fig. S2. Selection of samples for cytogenetics analysis.** (A) After the construction of copy number profiles with QDNAseq, we compared the measured (σ) and expected (Eσ) standard deviation of each sample to identify cases with very noisy profiles (red dots). (B) The distribution of the difference between σ and Eσ. Samples excluded from our statistics are highlighted in red. (C) The percentage of uninjected, Cas9 control and targeted samples with whole or sergmental losses/gains of chromosomes 5 and 7 according to their copy number profiles. (D) The percentage of uninjected, Cas9 control and targeted samples from blastocyst-stage embryos with whole or sergmental losses/gains of chromosome 6 according to their copy number profiles. The reported p-values are the result of two-tailed Fisher's tests.

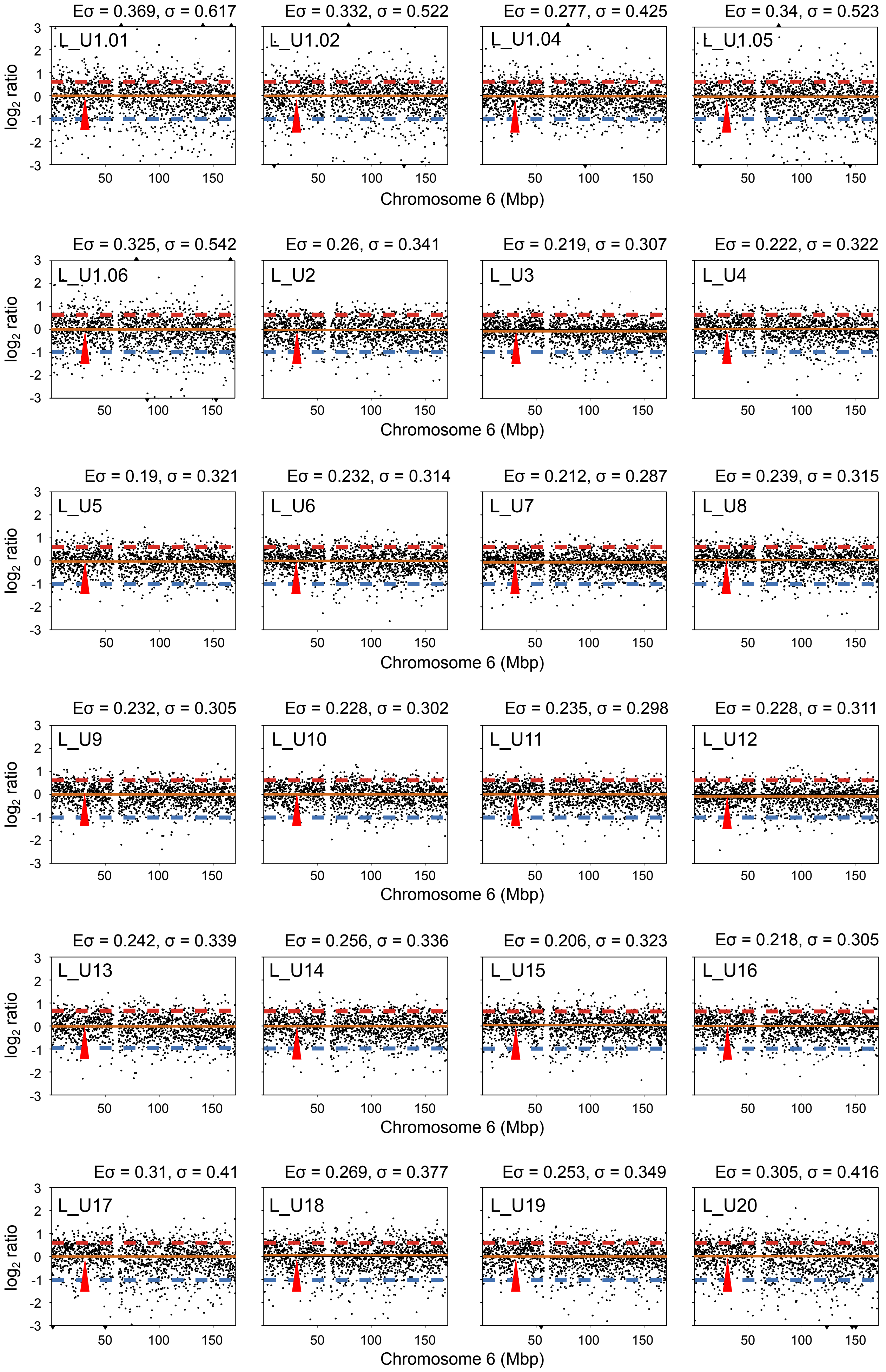


**Fig. S3. Zoomed-in view of the chromosome 6 copy number profile for all Uninjected samples with good quality low-pass WGS data.** Whole and segmental losses/gains of chromosome 6 have been highlighted. The approximate position of the *POU5F1* gene is indicated by a red arrow. The red dashed line indicates a copy ratio of 3:2, while the blue dashed line corresponds to a copy ratio of 1:2.

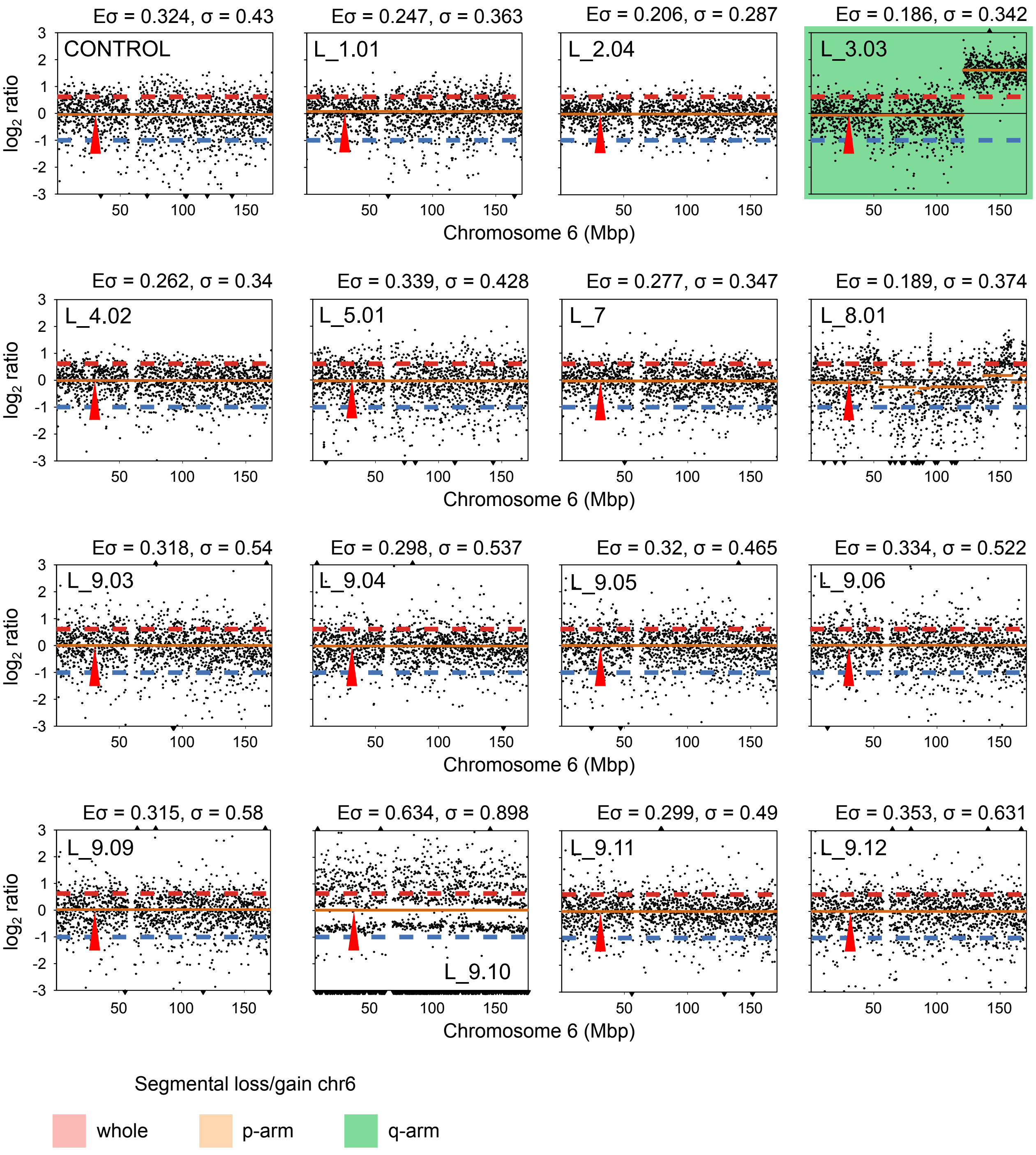


**Fig. S4. Zoomed-in view of the chromosome 6 copy number profile for all Cas9 control samples with good quality low-pass WGS data.** Whole and segmental losses/gains of chromosome 6 have been highlighted. The approximate position of the *POU5F1* gene is indicated by a red arrow. The red dashed line indicates a copy ratio of 3:2, while the blue dashed line corresponds to a copy ratio of 1:2.

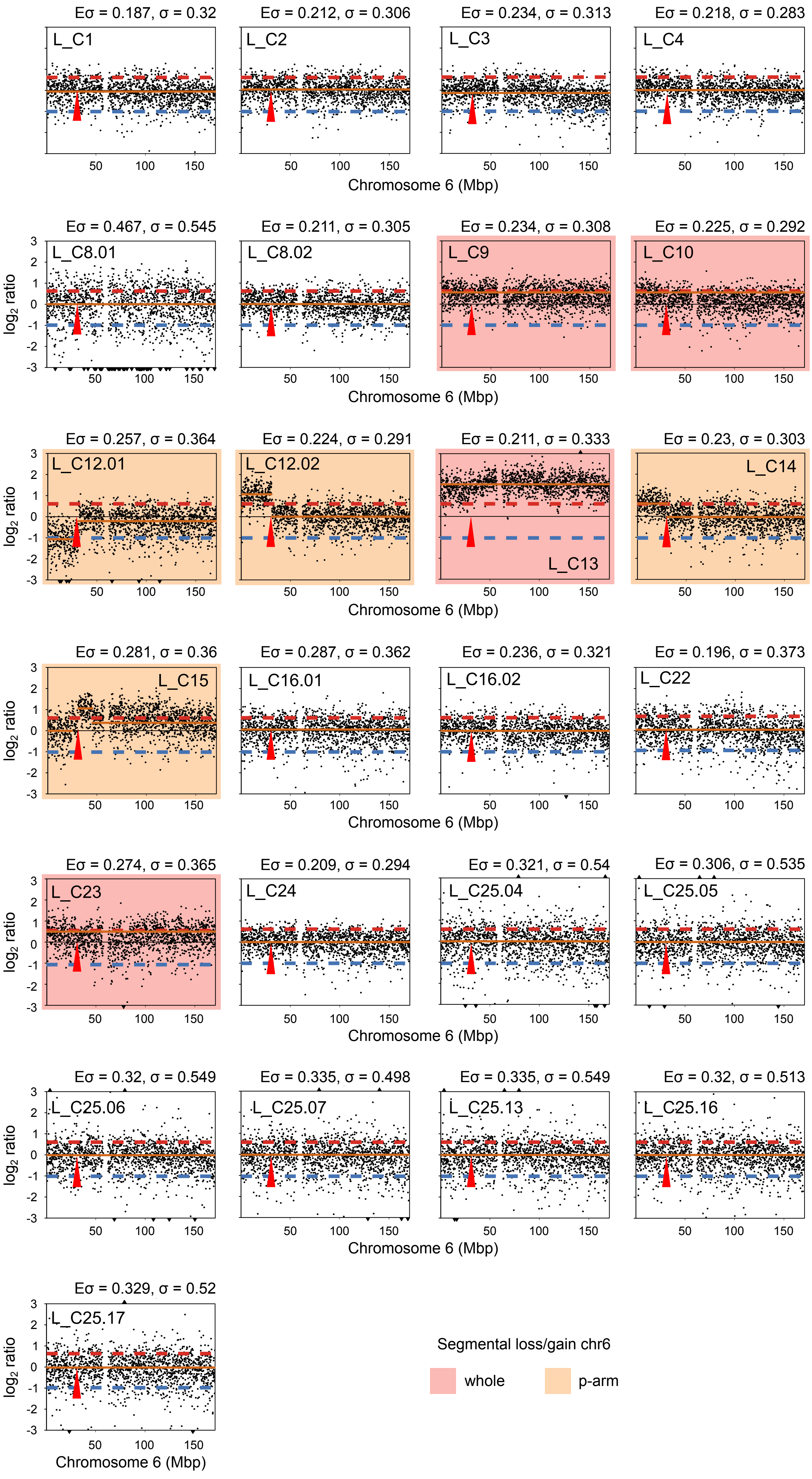


**Fig. S5. Zoomed-in view of the chromosome 6 copy number profile for all Targeted samples with good quality low-pass WGS data.** Whole and segmental losses/gains of chromosome 6 have been highlighted. The approximate position of the *POU5F1* gene is indicated by a red arrow. The red dashed line indicates a copy ratio of 3:2, while the blue dashed line corresponds to a copy ratio of 1:2.

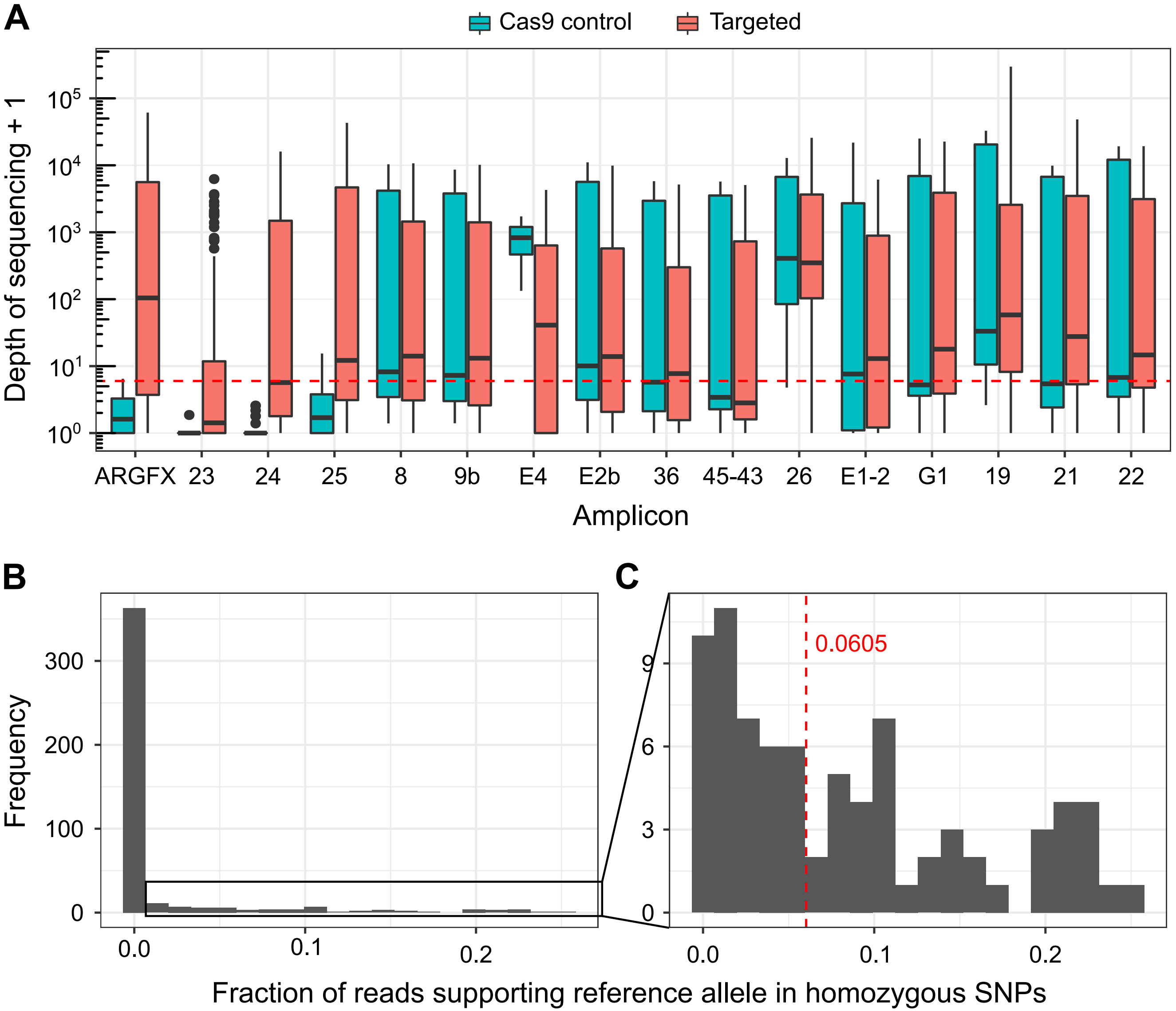


**Fig. S6. Considerations for SNP calling from deep sequencing data.** (A) The distribution of sequencing depths for the 16 PCR amplicons that we considered. The distributions for Cas9 control and OCT4-targeted samples are shown separately. The red dashed line indicates the minimum coverage of 5x that we required for SNP calling. (B) The distribution of the fraction of reads supporting the reference allele in homozygous SNP calls across all samples. (C) Zoomed-in view of the distribution for fractions above zero. The red dashed line corresponds to the median of the distribution and is the value that we used to control for the effect of allele over-amplification in our SNP calling pipeline.

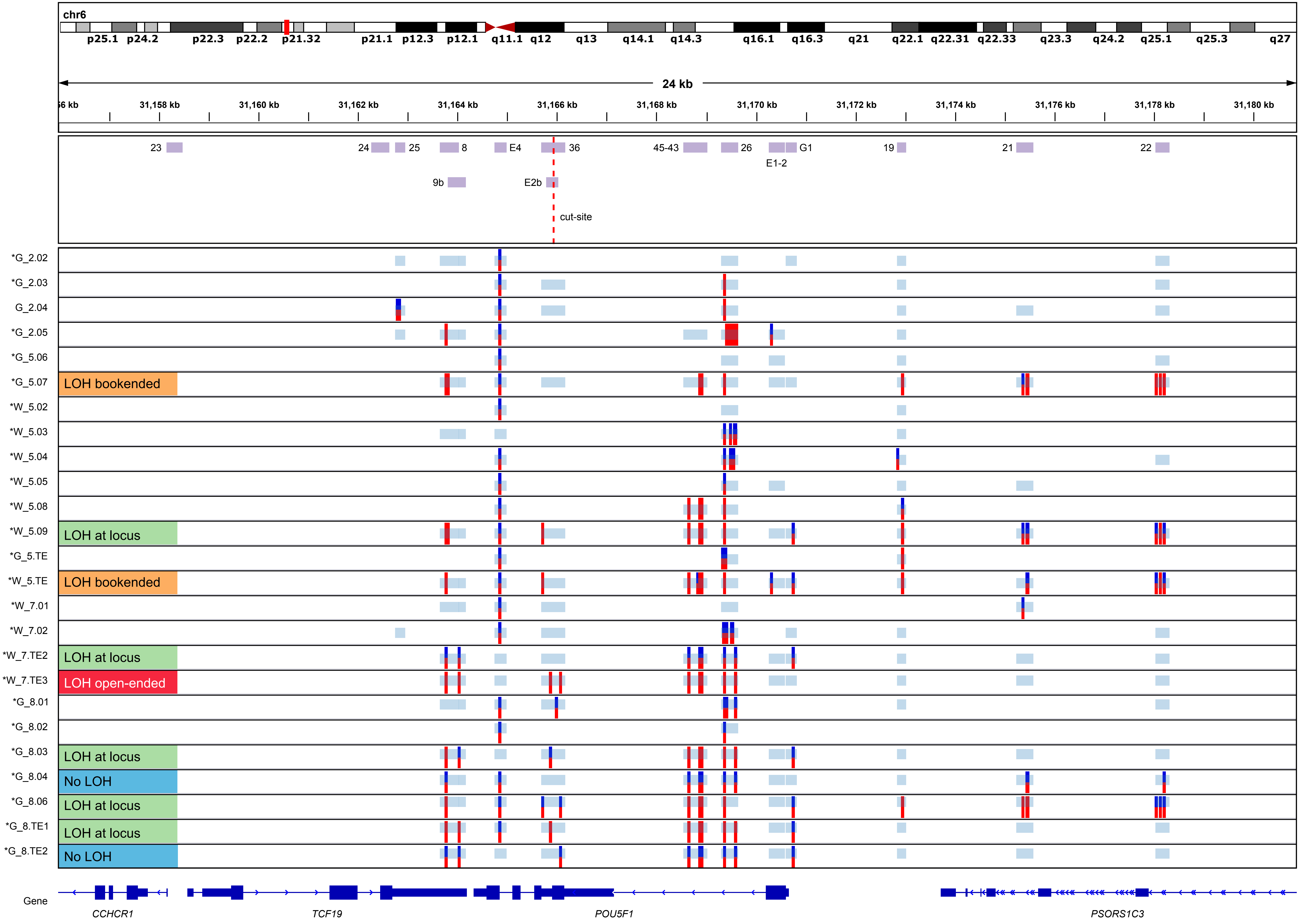


**Fig. S7. SNP profiles for Cas9 controls.** A type of loss-of-heterozygosity (LOH) event was assigned to samples with at least 10 amplicons sequenced at ≥ 5x depth of coverage. The *ARGFX* control amplicon (chr3:121,583,621-121,586,438) did not reach the 5x depth of coverage in samples marked with a *.

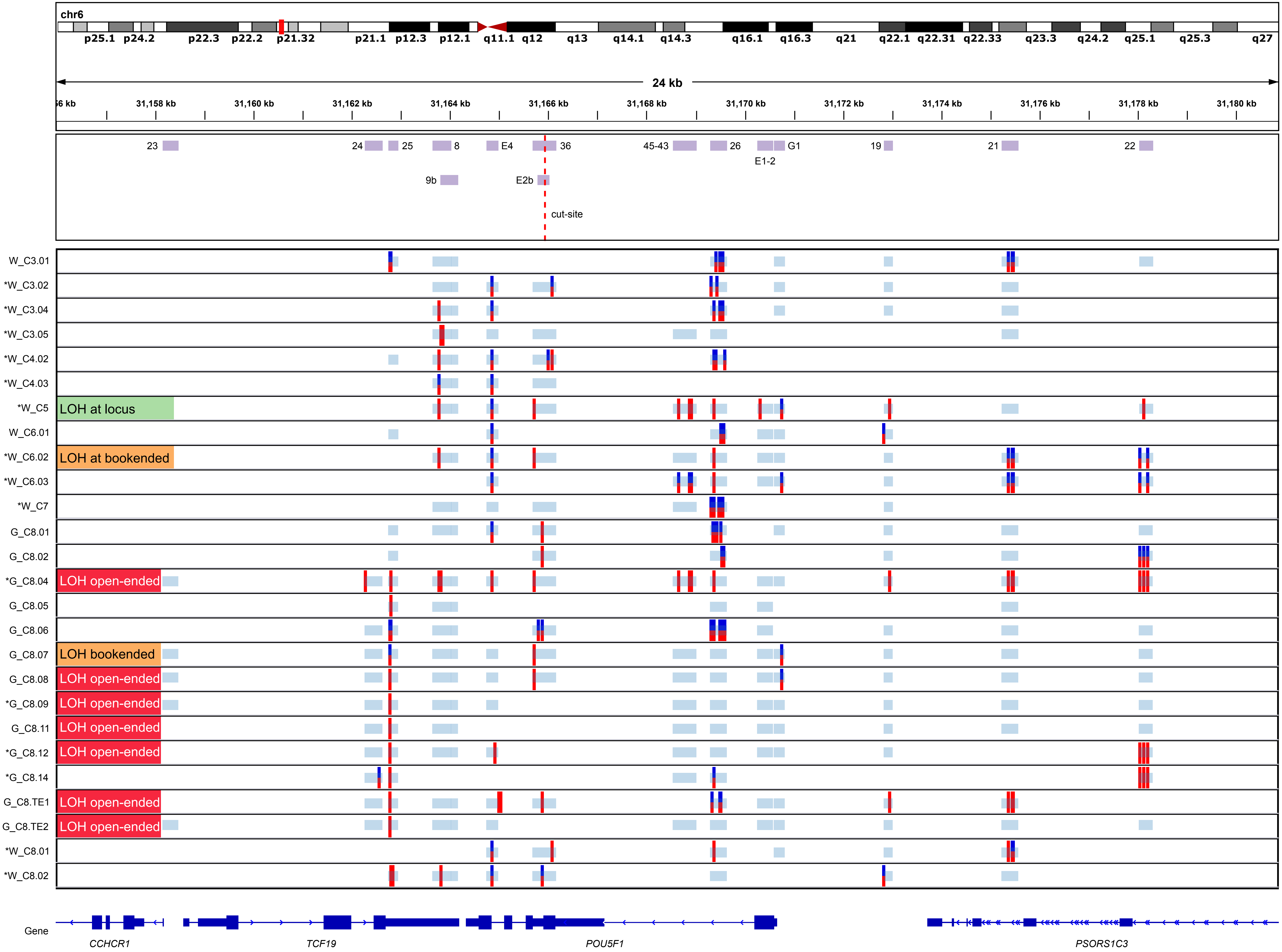


**Fig. S8. SNP profiles for single cells or trophectoderm biopsies from targeted embryos C3-C8.** A type of loss-of-heterozygosity (LOH) event was assigned to samples with at least 10 amplicons sequenced at ≥ 5x depth of coverage. The *ARGFX* control amplicon (chr3:121,583,621-121,586,438) did not reach the 5x depth of coverage in samples marked with a *.

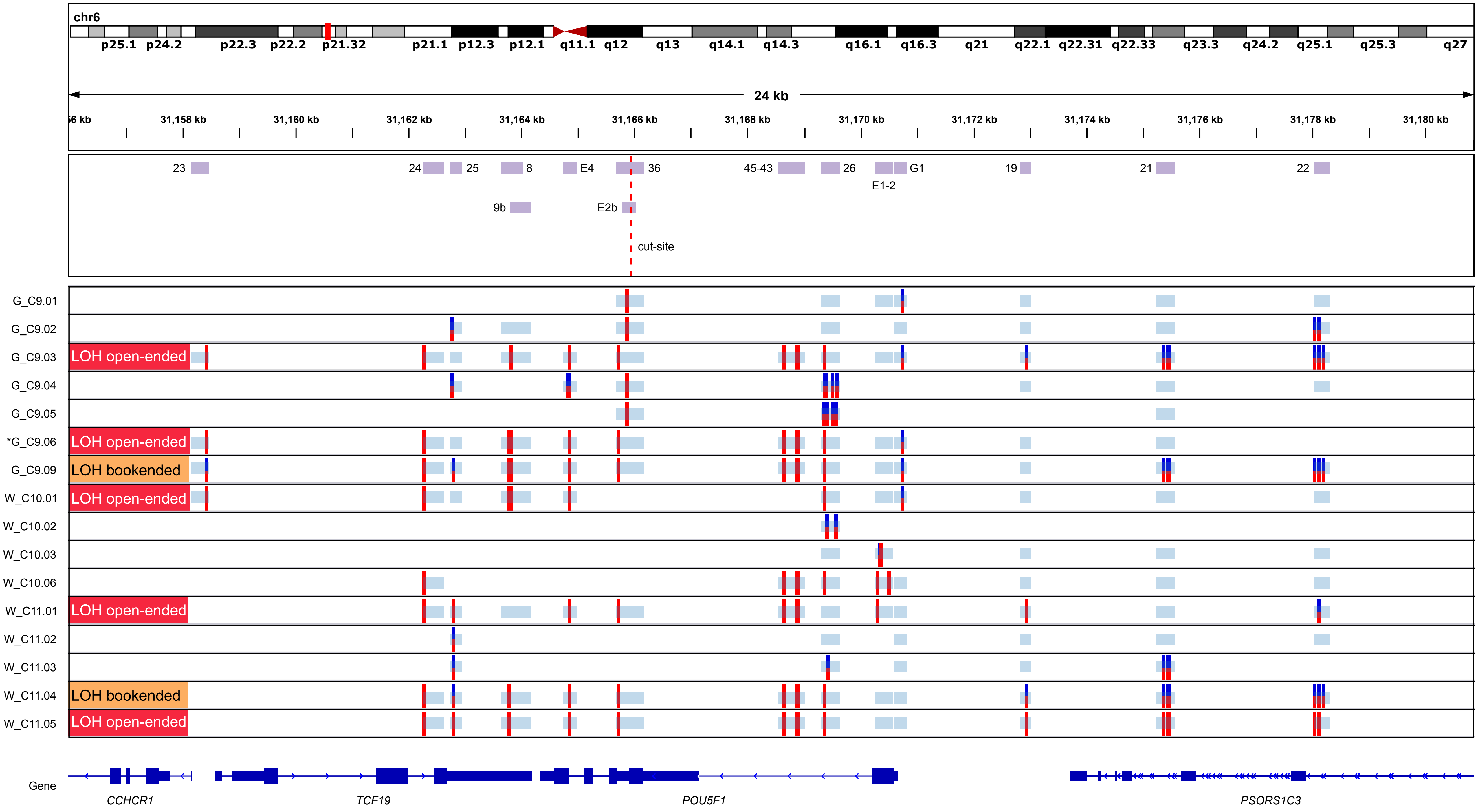


**Fig. S9. SNP profiles for single cells of targeted embryos C9-C11.** A type of loss-of-heterozygosity (LOH) event was assigned to samples with at least 10 amplicons sequenced at ≥ 5x depth of coverage. The *ARGFX* control amplicon (chr3:121,583,621-121,586,438) did not reach the 5x depth coverage in samples marked with a *.

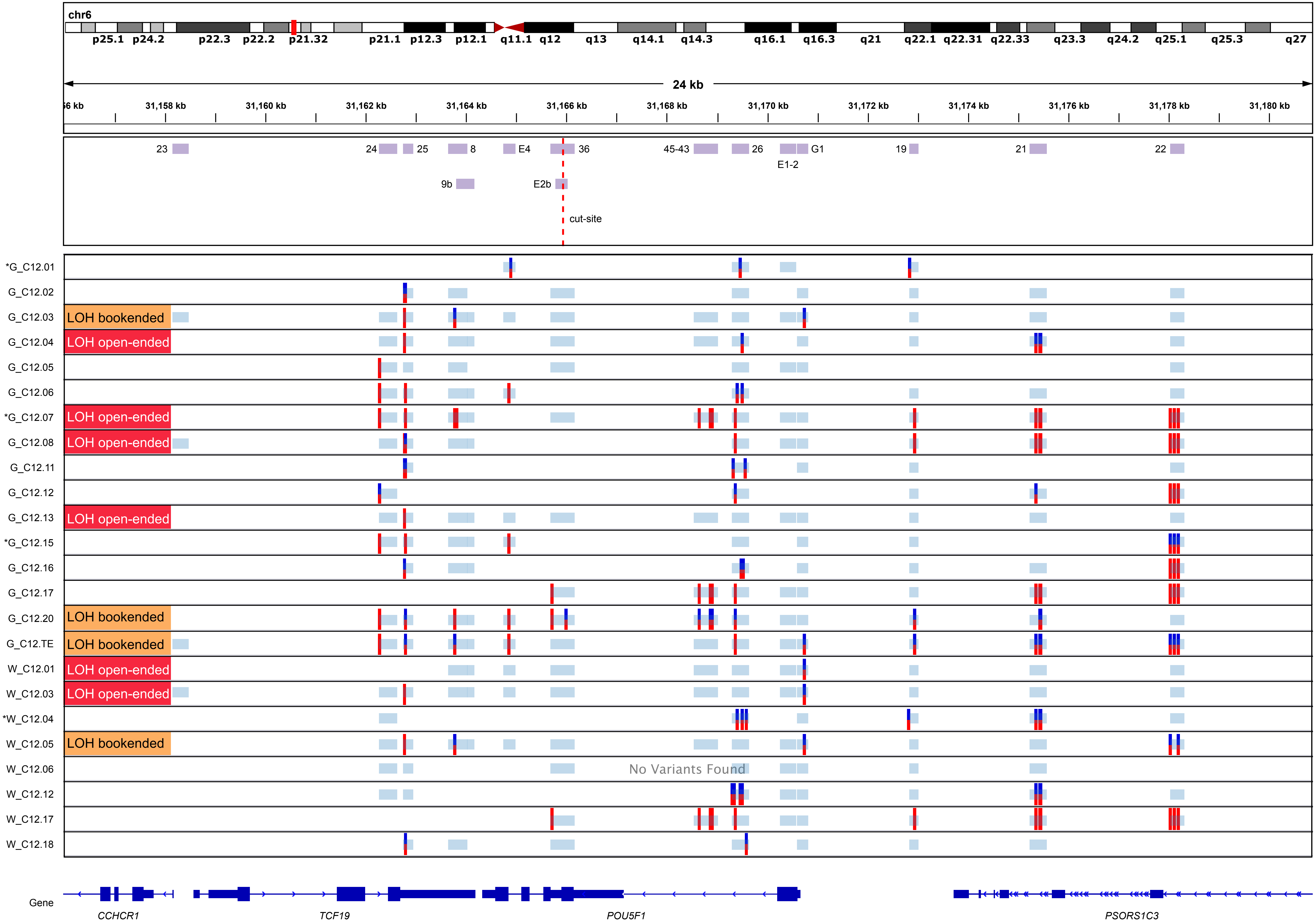


**Fig. S10. SNP profiles for single cells or trophectoderm biopsies from targeted embryos C12.** A type of loss-of-heterozygosity (LOH) event was assigned to samples with at least 10 amplicons sequenced at ≥ 5x depth of coverage. The *ARGFX* control amplicon (chr3:121,583,621-121,586,438) did not reach the 5x depth of coverage in samples marked with a *.

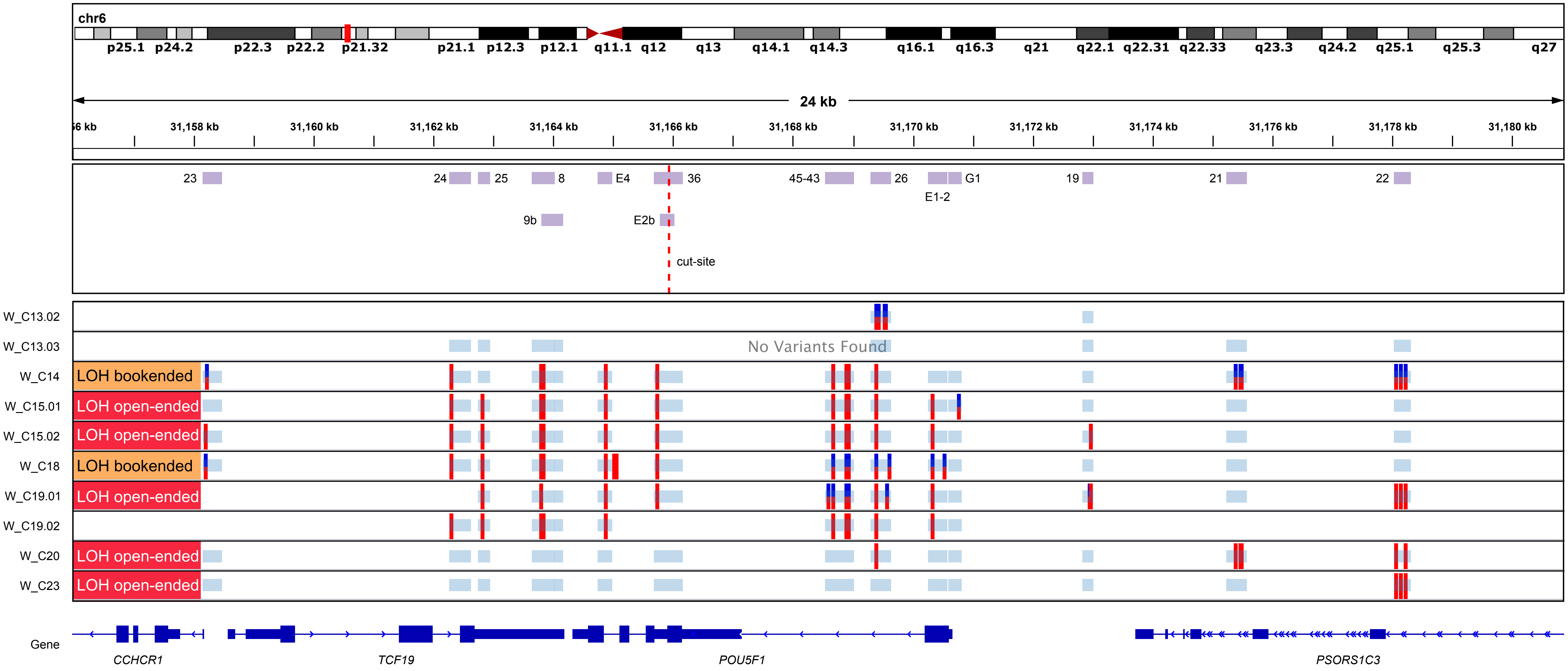


**Fig. S11. SNP profiles for single cells of targeted embryos C13-C15, C18-C20 and C23.** A type of loss-of-heterozygosity (LOH) event was assigned to samples with at least 10 amplicons sequenced at ≥ 5x depth of coverage. The *ARGFX* control amplicon (chr3:121,583,621-121,586,438) did not reach the 5x depth of coverage in samples marked with a *.

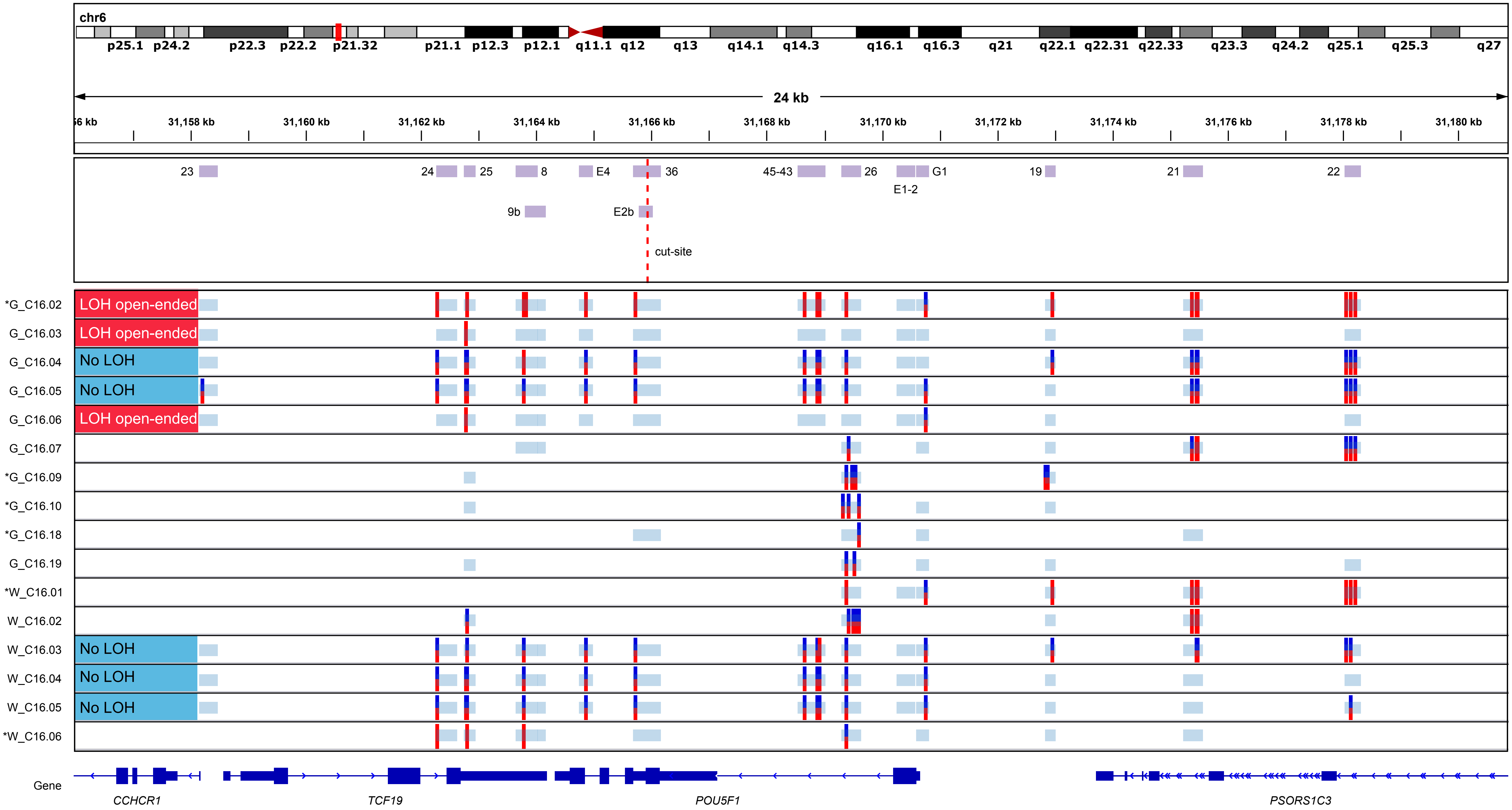


**Fig. S12. SNP profiles for single cells of targeted embryos C16.** A type of loss-of-heterozygosity (LOH) event was assigned to samples with at least 10 amplicons sequenced at ≥ 5x depth of coverage. The *ARGFX* control amplicon (chr3:121,583,621-121,586,438) did not reach the 5x depth of coverage in samples marked with a *.

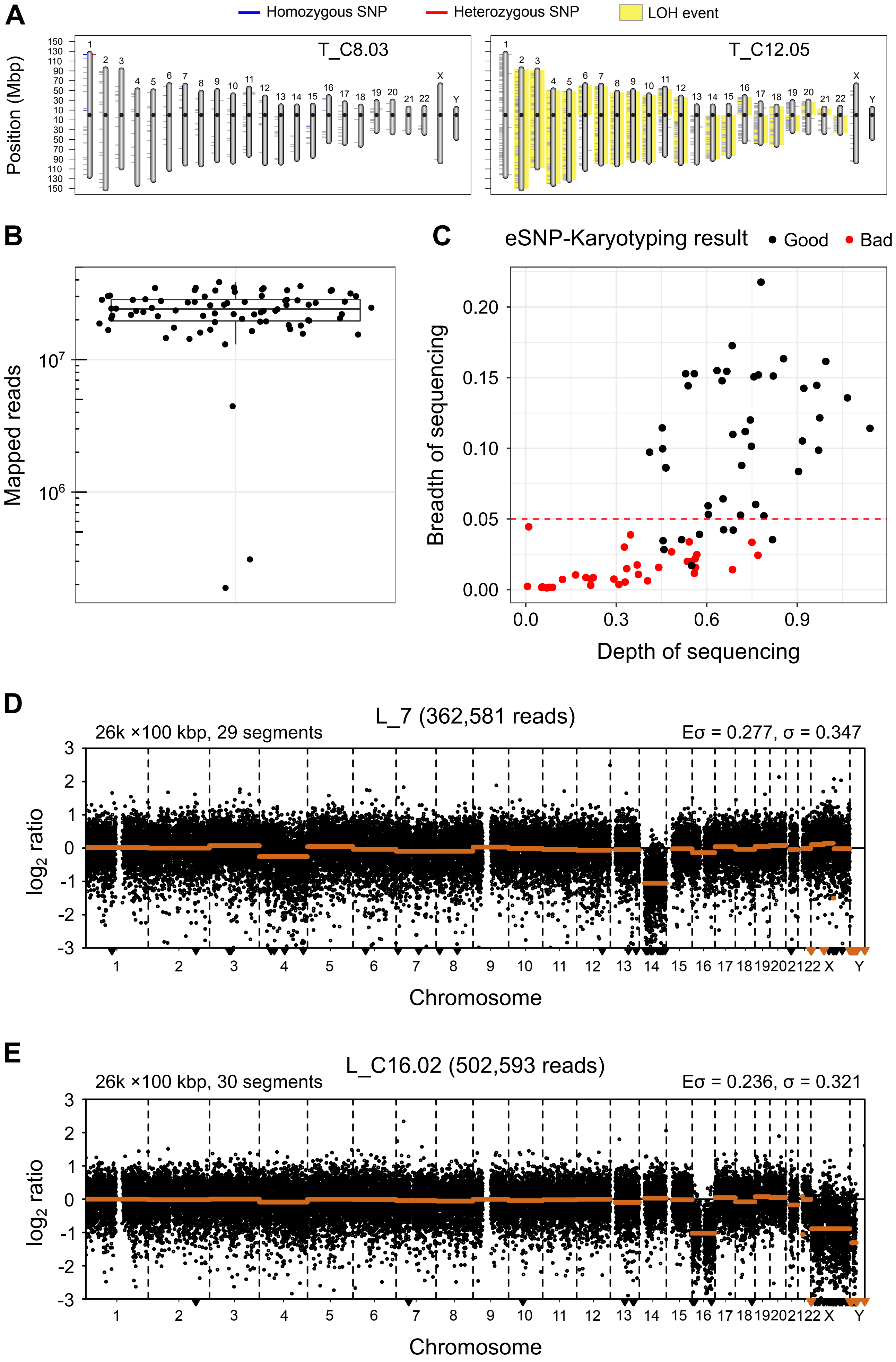


**Fig. S13. Selection of samples for transcriptome-based analyses.** (A) eSNP-karyotyping loss-of-heterozygosity (LOH) profiles for two low quality scRNA-seq samples. (B) Distribution of scRNA-seq mapped reads across samples. Except for three outliers, most samples have ~25 million mapped reads. (C) Breadth of sequencing is in better agreement with the quality of the eSNP-karyotyping results than the number of mapped reads (see B) and the depth of sequencing. Therefore, we used a conservative threshold of 0.05 breadth of sequencing (red dashed line) to select samples for transcriptome-based analyses (note that eight good quality samples were discarded under this cut off). (D) Copy number profile of sample L_7 that highlights the agreement between the low-pass WGS (loss of chromosomes 4 and 14) and the transcriptome-based karyotypes shown in Fig. 4A and B (sample T_7.01). (E) Copy number profile of sample L_C16.02 that highlights the agreement between the low-pass WGS (loss of chromosome 16) and the transcriptome-based karyotypes shown in Fig. 4A and B (sample T_C16.06).

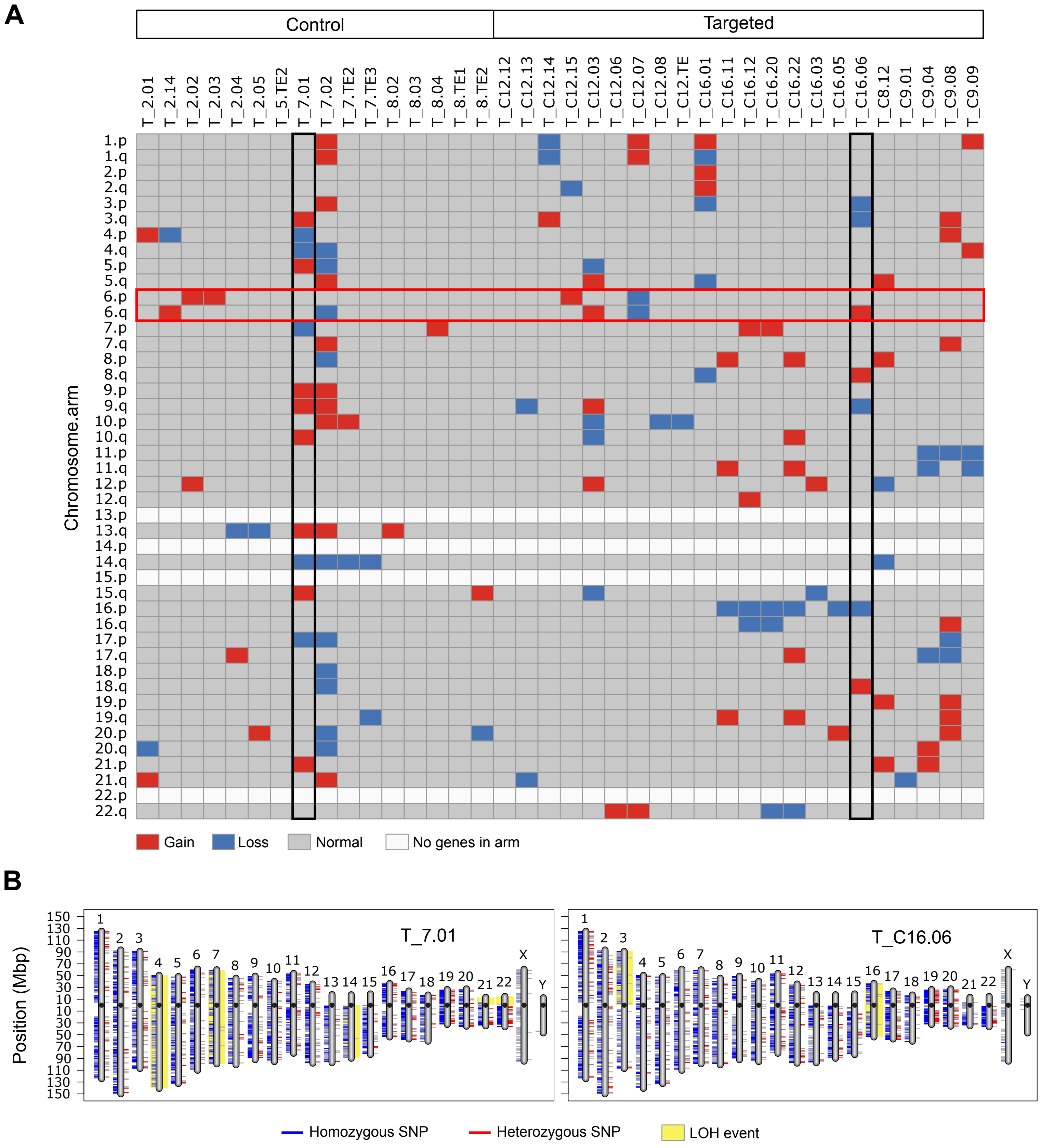


**Fig. S14. Transcriptome-based karyotypes.** (A) Digital karyotype based on the total gene expression deviation from the average of each chromosome arm (z-score-karyotyping). Chromosome 6 and two samples have been highlighted. (B) The loss-of-heterozygosity (LOH) profile of the two samples highlighted in A. These profiles were constructed with the eSNP-Karyotyping pipeline, which is also transcriptome-based. Note that the chromosome losses identified by this method were also captured by the karyotype in A.

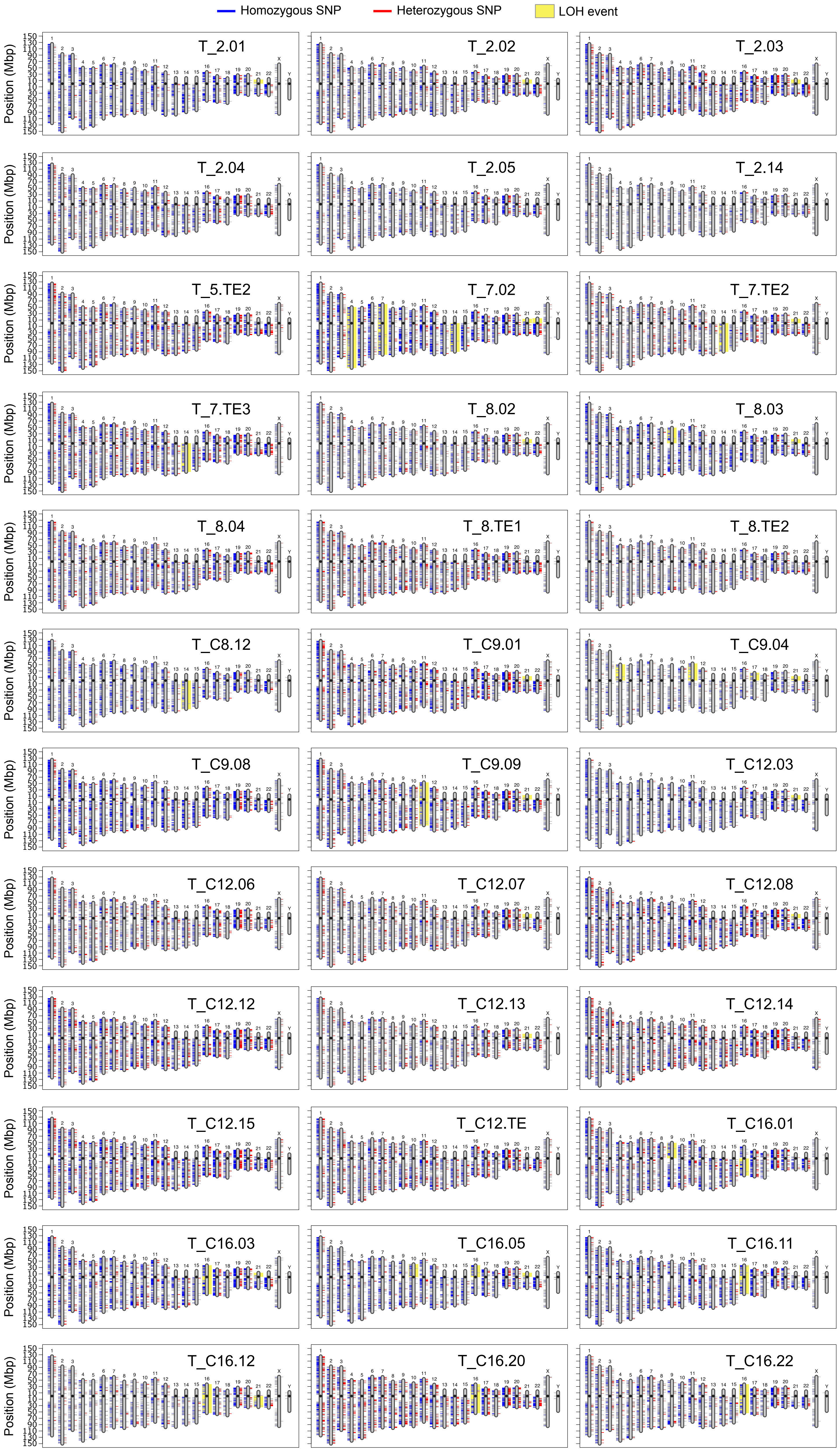


**Fig. S15. eSNP-Karyotyping results.** Loss-of-heterozygosity (LOH) profiles constructed with eSNP-Karyotyping for all samples with good quality scRNA-seq data.
